# Mechanical model of muscle contraction. 6. Calculations of the tension exerted by a skeletal fiber during a shortening staircase

**DOI:** 10.1101/2019.12.16.878876

**Authors:** Louvet S.

## Abstract

Accompanying Paper 1 tests a theoretical relationship between force and shortening velocity of a muscle fiber without justifying its validity. Paper 2 determines the kinematics and dynamics of a myosin II head during the working stroke (WS). Paper 3 imposes the Uniform law as a density representative of the orientation of the levers belonging to the WS heads. By support of these works, Papers 4 and 5 put into equation the evolution of the tension during the four phases of a length step. The present paper closes all six articles by imposing two tasks on itself. The first purpose is to apply the theoretical elements developed for a length step to a succession of identical length steps, otherwise known as shortening staircase. With the values of the geometric and temporal parameters assigned to a myosin head in Papers 1 to 5, a correct adjustment is established between the theoretical tension deduced from our model and the experimental tension published in 1997 by a team of Italian researchers relating to nine shortening staircases performed on the same fiber. In particular, we obtain the equation of the tension reached at the time end of the step (T*) which remains constant step by step as soon as the shortening of a half-sarcomere exceeds 17 nm. The second objective is to find and explain the equation of the Force-Velocity curve introduced *ex abrupto* into Paper 1: by decreasing the size and duration of the steps, the staircase tends towards a constant slope line corresponding to a continuous speed shortening. By applying the methods of infinitesimal calculus to the different formulations leading to T*, we deduce the Force-Velocity relationship (see Supplement S6.L). And the circle is complete.

## Introduction

An isolated muscle fiber is tetanized under isometric conditions until the tension reaches a maximum plateau (T0). Then, the fiber is shortened according to a series of steps with identical length (δX_stair_) and duration (τ_stair_). The temporal kinetics of this series of shortenings is similar to a succession of steps and consequently to a staircase. The tension (T_i_) is measured at the time end of step n° i with i integer. It is observed that T_i_ decreases regularly for the first steps and then remains constant [1,2,3,4] from a step numbered n*. For any step with an index greater than n*, the kinetics of the tension is an exact copy of that of the step n*, a temporal evolution that will be qualified as a "repetitive regime": the fiber is in an out-of-equilibrium thermodynamic state but reproducible from one step to the next. Figure 1 in [4] shows nine plots of relative tension (pT) as a function of time (t) for nine staircases singled out by different values of δX_stair_ and τ_stair_. Our first objective is to simulate the nine kinetics as closely as possible by basing on the time equations developed in accompanying Paper 5. The second purpose is to find the equations of the Force-Velocity relationship delivered to Paper 1 using the results established for the repetitive regime.

**Fig 1.**
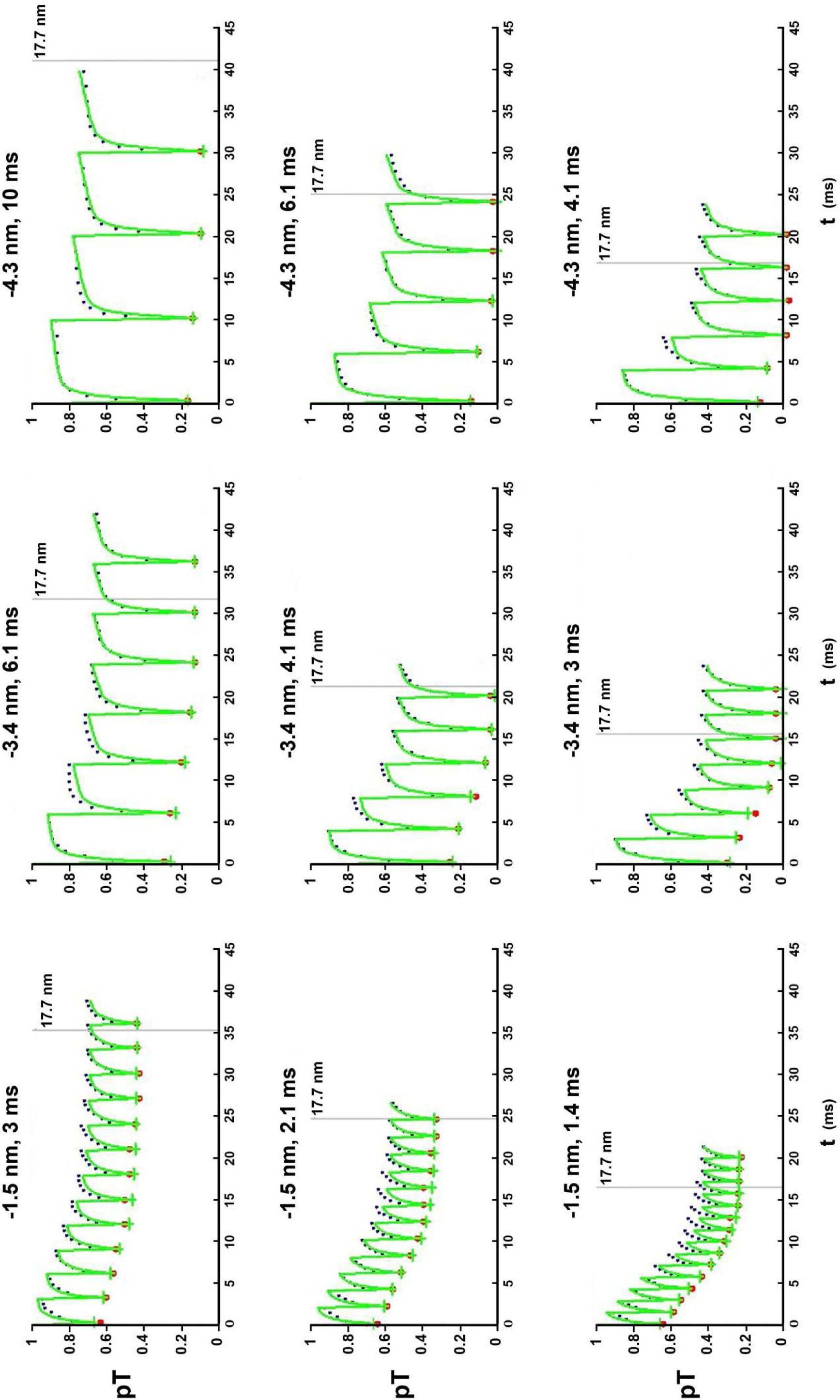
Time evolution of the theoretical tension (continuous green line) in response to a shortening staircase; the size and duration of the staircase step are indicated above each of the 9 graphs. The green crosses represent the tensions calculated at the end of phase 1 of each step (see explanations in the text). The black and red dots are from Fig 1 in [4]. The red dots represent the tensions collected at the end of phase 1 of each step.

## Methods

For staircase steps with an index i equal to or greater than n*, the tension T_i_ is a constant that is called T*.

During τ_stair_, each half-sarcomere (hs) of the fiber is shortened at constant velocity during phase 1 of the step and then maintained under isometric conditions. The events that occur during each step are described and equated in accompanying Papers 4 and 5. The δX_Max_ range is divided into intervals by successive steps of δX_stair_ length (Fig L1 of Supplement S6.L). In each interval, the contributions to the calculation of the tension related to the above events are determined in time and number of myosin heads involved. A step-by-step description from the isometric tetanus plateau up to the repetitive regime is given in paragraph L.3 of Supplement S6.L.

At step i of staircase k, the respective contributions to the calculation of the tension at both sides of a hs are summed using the master equation presented in Paper 5:

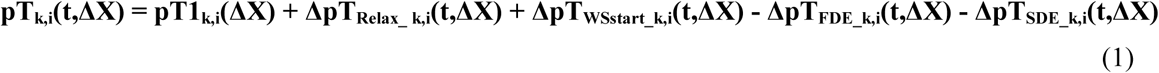

where **k** is the index number of the staircase, integer varying from 1 to 9

**i** is the index number of the staircase step, integer varying from 1 to 15, at most

**t** is the time such that:

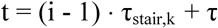

where τ_stair,k_ is the duration common to all steps of the staircase k; τ is the instantaneous time internal to the step i

**∆X** is the total shortening of a hs such that:

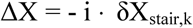

where δX_stair,k_ is the length common to all steps of the staircase k

**pT1_k,i_** is the value of relative tension at the end of phase 1 for step i of staircase k in the presence of viscosity

**∆pT_Relax_k,i_** is the positive contribution from the relaxation caused by the disappearance of viscous forces during phase 2 for step i

**∆pT_WSstart,k,i_** is the positive contribution of elastic origin coming from heads that initiate a WS in one of the 3 modes, fast, slow or very slow

**∆pT_FDE_k,i_** is the negative contribution caused by the rapid detachment, an event related to WS heads whose lever has an angle θ less than or equal to θ_down_

**∆pT_SDE_k,i_** is the negative contribution generated by the slow detachment, an event that only concerns WS heads whose lever has an angle θ between θ_down_ and θ_T_

### Calculation of the tension at the end of phase 1 for the step i of the staircase k

The relative tension at the end of phase 1 (pT1_k,i_) referred to the tension of the isometric tetanus plateau (T0_c_), a common value at the start of the nine staircases, is formulated according to equation (J64) given in sub-paragraph J.11.2 of Supplement S4.J to Paper 4:

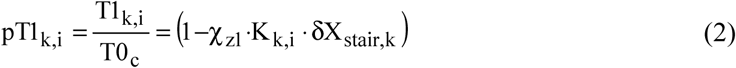

where k and i are the index numbers of the staircase and step concerned

χ_z1_ is the elastic origin slope in Zone 1 Enlarged, slope which we assume to be constant; K_k,i_ is the viscous origin multiplier coefficient formulated according to equality (J66):

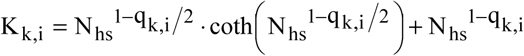

where q_k,i_ is the referent parameter of the presence of viscosity during phase 1 of step i of staircase k, whose equation is proposed in (J68):

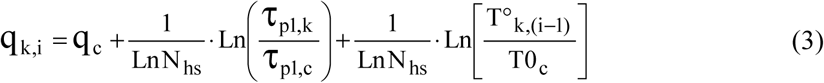

where q_c_ and τ_p1, c_ are the viscous parameter and the duration relative to phase 1 of the first step of one of the 9 staircases which serves as a reference; τ_p1,k_ is the average duration of phase 1 relative to phase 1 of all steps of staircase k; N_hs_ is the number of hs per myofibril; T°_k,(i-1)_ is the tension reached at the time end of the previous step with T°_k,0_ = T0_c_

For a correct application of formula (2) and (3), it is necessary that the step length is in Zone 1 Enlarged. According to equation (22) in Paper 4, the following condition is required:

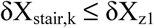

The δX_z1_ value (Table 2) implies:

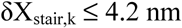

For staircases 7, 8 and 9, it will be assumed that the above condition is extended to δX_stair,k_ ≤ 4.3 nm.

## Algorithmic

Computer programs are written in Visual Basic 6. The calculations are developed in the form of algorithms to obtain the plots of the relative tension (pT) as a function of time (t) for a staircase, whatever the duration and length of the step if δX_stair_ ≤ 4.3 nm.

### Adequacy between theoretical and experimental kinetics

By allocating values to the model data that are compatible with those in the literature and those used in the calculations of Papers 1 to 5, nine pT kinetics are determined for nine different pairs of values from δX_stair_ and τ_stair_, identical pairs to those defining the nine staircases in Fig 1 in [4].

The adjustment between the nine theoretical kinetics from the model equations and the nine experimental kinetics in Fig 1 from [4] is done visually by the trial and error method.

## Results

### Application of the results of Papers 1 to 5 to a shortening staircase

The time equation (1) is represented by a green solid line on the 9 graphs in Fig 1, each relative tension plot being the controlled response to a shortening staircase. The values of the duration and length of a standard step, specific to each staircase, are displayed in Table 1 and are reported in the title of each corresponding graph. The average reference time for phase 1 of a step (τ_p1,c_) is taken relative to the staircase n° 6 (Table 1; light yellow column). The parameter values useful for constructing the nine plots are presented in Table 2. They are identical for all the steps of the nine staircases. There is an acceptable agreement between theoretical calculations (green line) and experimental measurements (black dots) taken from the nine curves in Fig 1 from [4].

**Table 1.** Data specific to each of the 9 staircases.

| k | 1 | 2 | 3 | 4 | 5 | 6 | 7 | 8 | 9 |
| --- | --- | --- | --- | --- | --- | --- | --- | --- | --- |
| $\delta X_{\text{stair}}$ | 1.5 nm | 1.5 nm | 1.5 nm | 3.4 nm | 3.4 nm | 3.4 nm | 4.3 nm | 4.3 nm | 4.3 nm |
| $\tau_{\text{stair}}$ | 3 ms | 2.1 ms | 1.4 ms | 6.1 ms | 4.1 ms | 3 ms | 10 ms | 6.1 ms | 4.1 ms |
| $\tau_{p1,k}$ | 0.16 ms | 0.15 ms | 0.15 ms | 0.18 ms | 0.17 ms | 0.2 ms | 0.26 ms | 0.24 ms | 0.23 ms |

**Table 2.** Data relating to a fiber isolated from the *tibialis anterior* muscle of *Rana Esculenta*. The values are identical for all stairs of the 9 staircases in Fig 1.

|  |  |
| --- | --- |
| $\Gamma$ | 4 °C |
| $T0_c$ | 200 kPa |
| $L0$ | 4.58 mm |
| $N_{hs}$ | 4400 |
| $L0_s$ | 2.1 $\mu\text{m}$ |
| $P_{\text{startF}}$ | 0.6 |
| $P_{\text{startS}}$ | 0.3 |
| $\delta X_{\text{Max}}$ | 11.7 nm |
| $\delta X_T$ | 8 nm |
| $\delta X_{\text{pre}}$ | 6 nm |
| $\delta X_{z1}$ | 4.2 nm |
| $q_c$ | 1.91 |
| $\tau_{p1,c}$ | 0.2 ms |
| $\tau_{\text{preSB}}$ | 1.5 ms |
| $\tau_{\text{SB}}$ | 4 ms |
| $\tau_{\text{Relax}}$ | 0.5 ms |
| $\tau_{\text{startF}}$ | 0.7 ms |
| $\tau_{\text{preS}}$ | 3 ms |
| $\tau_{\text{startS}}$ | 30 ms |
| $\tau_{\text{preVS}}$ | 80 ms |
| $\tau_{\text{startVS}}$ | 90 ms |
| $\tau_{\text{preFDE}}$ | 1 ms |
| $\tau_{\text{FDE}}$ | 5 ms |
| $\tau_{\text{preSDE}}$ | 5 ms |
| $\tau_{\text{SDE}}$ | 18 ms |

The relative tensions at the end of phase 1 of each staircase step (pT1_k,i_) are determined using equations (2) and (3) and appear with a green cross (Fig 1); these theoretical values adjust to the experimental points represented by a red point. A correction to the value of q_c_ is made for the three staircases with a length step equal to 1.5 nm: the value of q_c_ is kept for the first step but gradually decreases to the value of 1.81 which becomes constant from the fifth step. A possible explanation for this empirical correction is a more important relaxation time for small length steps that reinforce the presence of viscosity. This phenomenon is also observed in Figs 1 and 3 of Paper 5 where the time constant (τ_Relax_) is equal to 0.5 ms for length steps equal to or less than 1.5 nm and decreases to 0.3 ms for steps greater than 1.5 nm.

### Repetitive regime

The repetitive regime occurs at step n* when all the "prepared" heads in a strong binding state and all the heads in a WS state initially at t=0 are finally detached. So from equality (L4) of Supplement S6.L transformed linearly according to (L1), it is checked approximately:

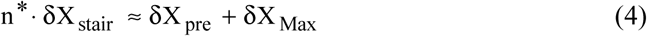

The data in Table 2 give:

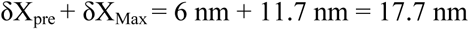

On each of the 9 graphs, this value is represented by a fine vertical line at the time of the corresponding shortening.

### Establishing the Force-velocity Equation

It is recalled that the shortening velocity of the fiber and its myofibrils (V) is related to the hs shortening velocity (u) as:

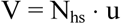

#### 1/ Event {WSstart}

A myosin head initiates a WS with the global event {WSstart} which is broken down into three separate events, {startF}, {startS} or {startVS}, when the initiation is fast, slow or very slow, respectively. Paragraph L.4 of Supplement S6.L is devoted to the calculation of the tension generated by the WS heads in repetitive regime. The result is formulated in (L15) and detailed with equations (L16a), (L16b) and (L16c) related to the initiation to the WS state according to the three modes.

In paragraph L.5, the two variables characterizing a staircase step, length (δX_stair_) and duration (τ_stair_), are gradually reduced until they tend towards infinitesimal values. This brings closer to a shortening performed at a constant speed (u) for each hs. The principles of differential and integral calculus are applied to the three expressions (L16a), (L16b) and (L16c) and provide the three components of the tension generated by the WS heads when a hs is shortening at steady velocity. The three positive contributions are formulated in (L24), (L26) and (L28) according to the three modes, fast, slow and very slow, respectively.

#### 2/ Event {SlowDE}

A myosin head slowly detaches according to the event {SlowDE}. Paragraph L.6 is devoted to the calculation of the tension generated by the slowly detaching heads in repetitive scheme. The result is formulated in (L39) and detailed with the six equations (L40a) to (L40f).

In paragraph L.7, the principles of infinitesimal calculation are applied to expressions (L40a) to (L40f) and provide the tension generated by slowly detaching heads when a hs is shortening at constant velocity (u). The six negative contributions are formulated from (L44a) to (L44f). Since the 3 components related to Type 2 of {SlowDE} expressed in (L44d), (L44e) and (L44f) are negligible, they are not included in the final equations to paragraph L.8 and Paper 1 for simplification purposes.

#### 3/ Accounting balance sheet

In paragraph L.8, the tension exerted at the endpoints of a myofibril shortening at a constant velocity (V) is calculated by summing the three contributions relating to the events {startF}, {startS} and {startVS}, and subtracting the contribution relating to the event {SlowDE}. The result is finalized with equation (L47) where the hs shortening velocity is expressed algebraically (u<0). Equation (L47) is divided into 4 equations in accompanying Paper 1, noted (7a), (8a), (9a) and (10a) where u is expressed in modulus.

The density (f) of the angle θ of the levers belonging to the WS heads during shortening at steady velocity is the algebraic sum of four densities:

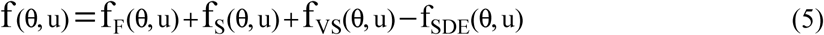

where f_F_, f_S_ and f_VS_ are the densities formulated in (L52a), (L52b) and (L52c), relative to an WS initiation fast, slow or very slow, respectively; f_SDE_ is the density formulated in (L53) relative to the slow detachment.

The density established in (5) is represented as a function of the angle θ by 9 curves according to 9 values of u (Fig 2). The data are from the "REF" column of Table 1 of Paper 1 where τ_preSB_ = 2.6 ms and τ_SB_ = 6 ms. The graph complies with the geometric criteria of a hs on the right. For a hs on the left, the signs of θ and u being opposite, the density plots are symmetrical with respect to the y-axis.

**Fig 2.**
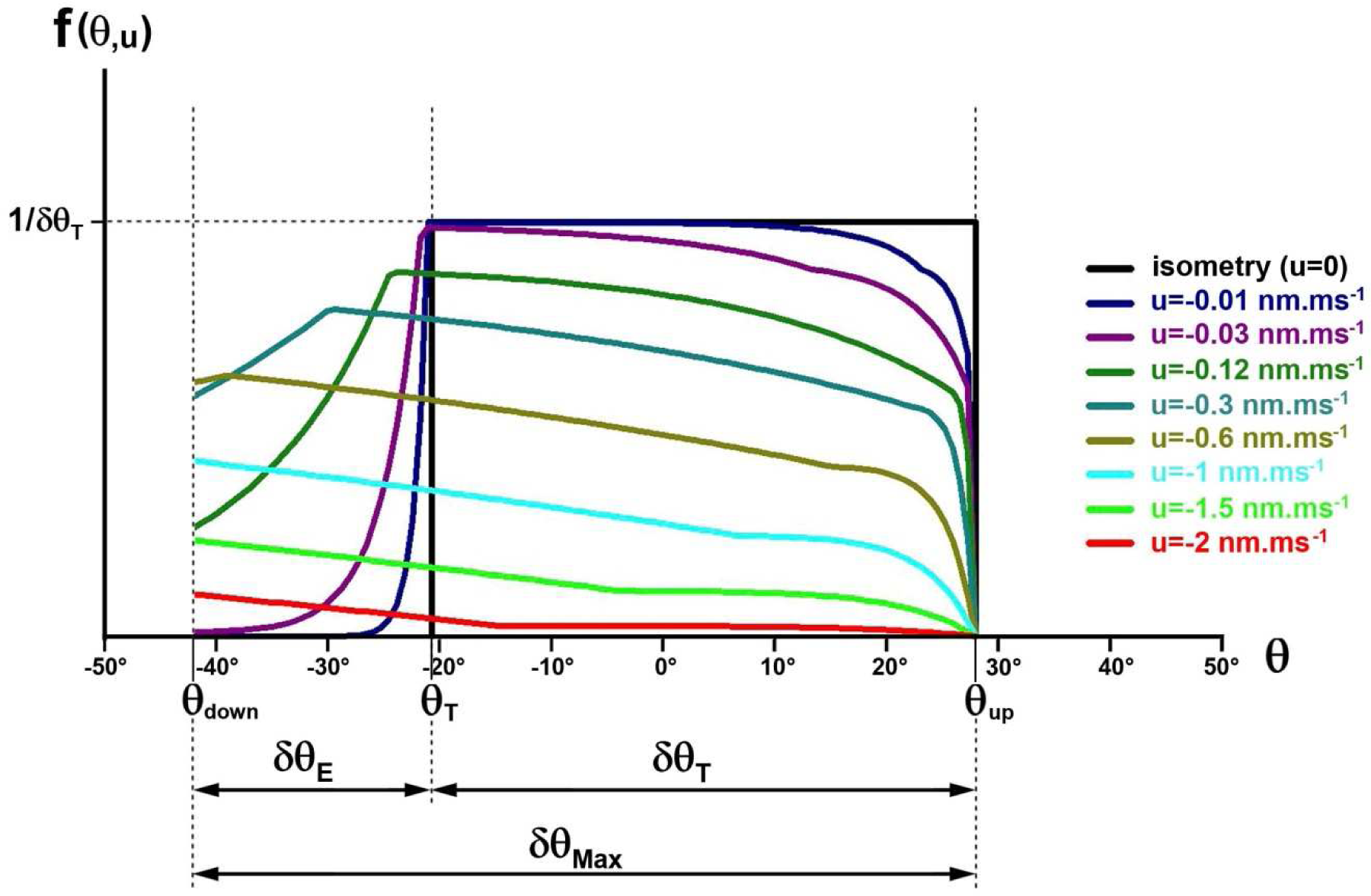
Density of the θ angle between θ_up_ and θ_down_ for 9 values of the shortening velocity (u) relative to a half-sarcomere on the right according to equation (5).

It is checked that if u tends towards 0, then f(θ,0) converges towards the Uniform law of δθ_T_ range, probability density characteristic of the isometric tetanus plateau (see Papers 3 and 4) represented by a black horizontal line between θ_up_ and θ_T_ (Fig 2).

## Discussion

### Validation of model equations

The theoretical and experimental tension kinetics for the 9 cases of examples in Fig 1 are consistent during a shortening staircase. The presence of the viscosity during phase 1 and the relaxation induced during phase 2 explain the kinetics of the first 3 milliseconds of each staircase step. The repetitive regime occurs when the myosin heads in Strong Binding "prepared" state and in Working Stroke state before the beginning of the staircase shortening have all detached, i.e. at the advent of step n* when the total hs shortening exceeds the length (δX_Max_+ δX_pre_), equal to 17.7 nm according to our calculations applied to the 9 staircases of Fig 1. By examining Fig L1f and L1g to Supplement S6.L, we note that densities f(θ) are close and that consequently the tension (T_n*-1_) at the time end of the preceding step n* is almost equal to T*; this explains why the repetitive regime can be empirically detected as soon as the index step (n*-1) is performed and why the value of 15 nm, lower than 17.7 nm, is given in [4] as the reference length for characterizing the repetitive regime. In all cases brought to our attention [2,3,4], the value of T_i_ becomes constant after an integer number of staircase steps whose sum of lengths is greater than 17 nm.

### Relationship between force and velocity of shortening

Based on the classical geometry of a myosin head and probabilistic criteria derived from the four events preparing and closing the working stroke state during the cross-bridge cycle, the developments of the Supplement S6.L lead to the Force-Velocity relationship exploited in Paper 1. It is reminded that for medium and high speeds, a corrective factor induced by the presence of viscosity must be taken into account in the calculations, decreasing the value of the tension provided in (L47).

### Density of the angle θ depending on the shortening speed u

Among the classical laws, the one that best approaches each of the nine distributions of the θ angle plotted on Fig 2 is the Uniform law. This finding supports the conclusions of H.E. Huxley and coauthors; see Table 1 and Fig 14 from [5]. This result also validates the equation (J80) of Supplement S4.J to Paper 4 where the instantaneous angular range δθ_T,i_, assumed to be uniform, increases from δθ_T_ to δθ_Max_ as the force step decreases; see Fig 6a of Paper 4 and the insets in Fig J14a and J14b from Supplement S4.J.

The rise of the intermediate angular range δθ_T,i_ from δθ_T_ to δθ_Max_ explains the apparent increase in the maximum WS step or stroke size of a myosin head from δX_T_ to δX_Max_ when the tension decreases (and the velocity enhances), this has already been mentioned in Discussion section of Papers 1 and 4.

### Homogeneity of data

The parameter values displayed in the tables of Papers 1 to 6 are similar. This comment is valid, on the one hand, for geometric data specific to a myosin head and hs of vertebrate and, on the other hand, for temporal variables relating to cross-bridge cycle reactions, with the exception of the time constants τ_preSB_ and τ_SB_, the study of which is covered in the following subsection. In particular, the standard values of the proportions of WS initiation in fast (p_startF_ ≈ 60% ±5%) and slow (p_startS_ ≈ 30% ±5%) mode are almost constant. These values are found independently for phase 4 of a force step (Paper 1), for phases 2, 3 or 4 of a length step (Paper 5) and for a staircase shortening (Paper 6). A geometric explanation has been suggested in the "Strong Binding Event" subsection of the Discussion section to Paper 5.

### Strong Binding state

This state characterizes the establishment of a strong binding (SB) between the myosin head and the actin molecule before the possibility of a rapid transition to the WS state through the {startF} event. The realisation of the {SB} event mentioned in paragraph L.2 is dependent on two time parameters τ_preSB_ and τ_SB_. The increase in one or both parameters results in a decrease in the proportion of the number of heads likely to initiate a WS quickly and implies a reduction of the tension. This decline is observed experimentally. An average velocity of shortening staircase (u*) is interpreted as the ratio of the length of the step to its duration, that is:

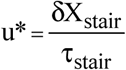

The tension (T*) at the time end of a staircase step in repetitive regime is always higher than the tension corresponding to the force step where each hs shortens at a steady velocity (u) equal to u*; see Fig 7 in [3] and Fig 3 in [4]. Thus, under identical experimental conditions, the values of the parameters τ_preSB_ and τ_SB_ determined for a shortening performed at steady velocity during phase 4 of a force step (Paper 1; Table 1) are amplified by about 60% compared to those associated with a staircase shortening (Paper 6; Table 1). A possible theoretical explanation is the transition to continuous.

This decrease in tension associated with the increase in τ_preSB_ and τ_SB_ is observed in Paper 1 for exogenous factors such as temperature (Tables 2 and Fig 4) and the tonicity of the Ringer solution (Table 3 and Fig 5).

It remains to be understood how the Strong Binding state prior to the rapid transition to the WS state operates. Should one or more intermediate steps and therefore one or more additional states be explored within the Strong Binding state as in the case of the transitional phases of the recovery stroke [6, 7]?

## Conclusion

Starting from the classical geometry of a myosin head, using probabilistic criteria derived from events of the cross-bridge cycle, and assuming the presence of viscosity, the model explains and predicts the evolution of the tension exerted at the two ends of a fiber disturbed by a length step (∆L<0) or by a force step (∆T<0).

The model also interprets the stretching of a fiber and the influence of temperature, which are the subjects of two future articles. For elongations (∆L>0), the bases are laid with the definition of Zone O in paragraph J.5 of Supplement S4.J to Paper 4. A first outline of the role of experimental temperature is given in Supplement S3.H of Paper 3 associated with sub-paragraph J.16.4 of Supplement S4.J.

## Supporting information

Data relating to computer programs used for Paper 6.

. Computer Programs for calculating and plotting kinetics of the nine shortening staircases

Supplementary Chapter. Theoretical study of a staircase shortening

## Acknowledgements

I thank Professor G. Piazzesi for allowing me to reproduce the points presented in Fig 1.

## Additional information

**S6.L Supplementary Chapter. Theoretical study of a staircase shortening.**

L.1 Step n° i belonging to the linear domain

L.2 Instantaneous proportion of heads that can quickly initiate a WS (recall)

L.3 Step-by-step description of a staircase shortening

L.4 Calculation of the tension generated by the WS heads in repetitive mode

L.5 Calculation of the tension generated by the WS heads when a hs is shortening at steady velocity u

L.6 Calculation of the tension generated by the slowly detaching heads in repetitive mode

L.7 Calculation of the tension generated by the slowly detaching heads when a hs is shortening at constant speed

L.8 Total with the 4 events {startF}, {startS}, {startVS} and {SlowDE}

L.9 Density of the θ angle as a function of the hs shortening speed (u)

**CP6 Supplementary Material. Computer Programs for calculating and plotting kinetics of the nine shortening staircases.** Algorithms are written in Visual Basic 6.

**DA6 Supplementary Material. Data relating to computer programs used for Paper 6.** Access Tables are transferred to Excel sheets.

