## Supplementary material for "Mechanical model of muscle contraction. 6. Calculations of the tension exerted by a skeletal fiber during a shortening staircase": . Computer Programs for calculating and plotting kinetics of the nine shortening staircases

### CP6 Computer Programs for calculating and plotting kinetics of the nine shortening staircases in Paper 6

The curves on the computer screen are done by calling the "SUB AAPH_ESC_trace()" routine.

Once the data have been recovered (See values displayed in the Table 1 and 2 of Paper 6 and in the Excel sheets of Supplement DA6, the experimental tension from Linari 1997article and the theoretical tension curves from model equations are plotted as a function of the time (Fig CP6.1).

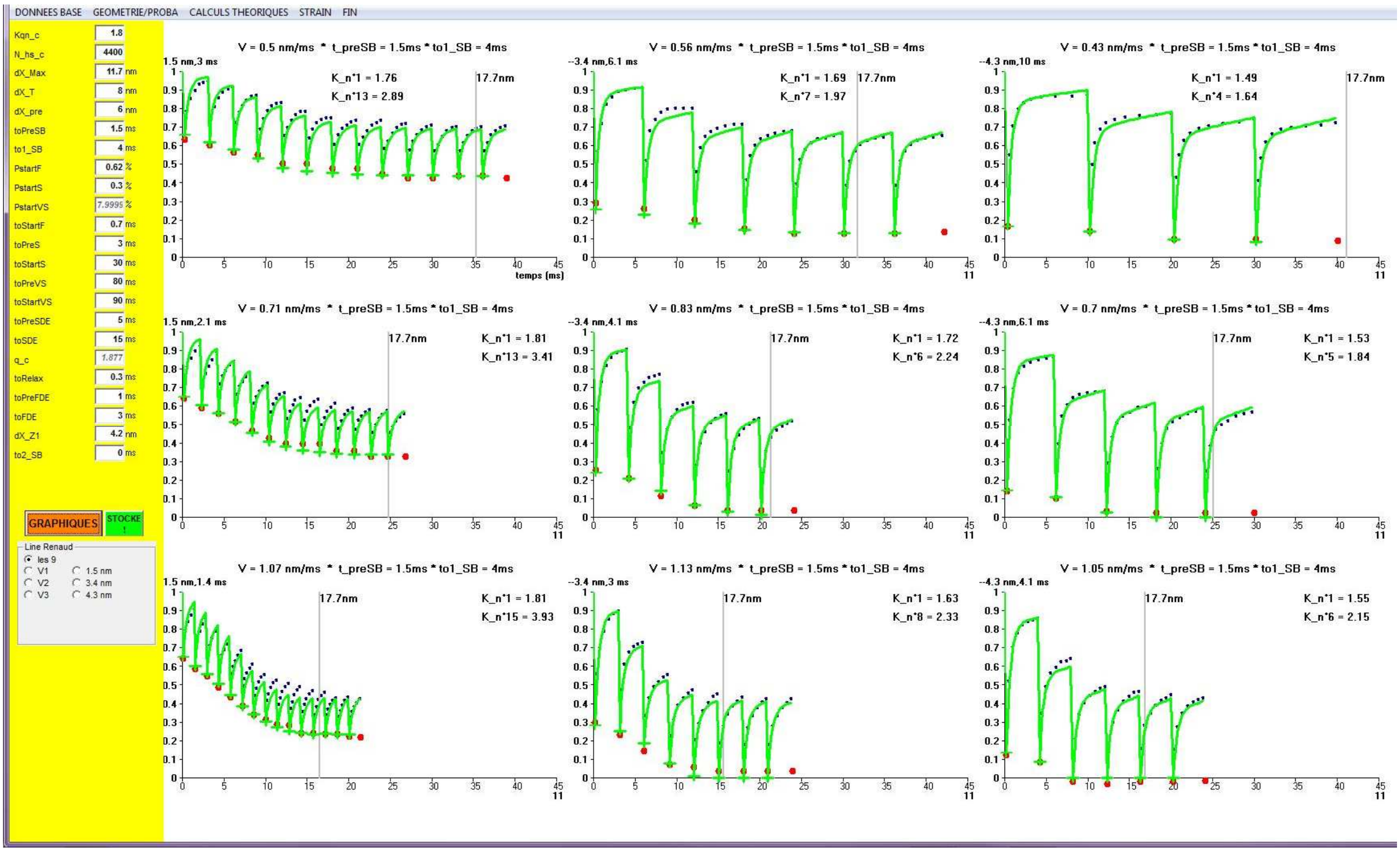


**Fig CP6.1 Screenshot after starting the the "Sub AAPH_ESC_trace()" routine.**

#### Sub AAPH_ESC_trace() '------ESC linari Ti T1

```
' choix
 'txV = "to_SB"
 txV = "t_preSB"
'principe :
' on travaille avec table ESC' je laisse tomber table _REMONTEE
If ressort.Tag <> "ESC" Then Stop
If mmm = 1 Then O.Cls
Np4 = 20 'nre de pts par step

'DONNEES DE BASE'--------------------
KqnZ1 = Val(TAM(0).Text) 'ne sert à rien
N_hs = Val(TAM(1).Text) 'ne sert à rien

L_WS = Val(TAM(2).Text) 'dX_Max = 12 nm

dX_T = Val(TAM(3).Text) 'dX_T =8.5 nm
aX = dX_T

L_preWS = Val(TAM(4).Text) 'dX_pre = 5.5 nm

to1_df = Val(TAM(5).Text) 't_pre_SB1 ms
to1 = Val(TAM(6).Text) '=to1_SB mais en fait varien en fonction de dX_step (voir +bas) ms
to2_SB = Val(TAM(22).Text)

P_startF = Val(TAM(7).Text)
P_startS = Val(TAM(8).Text)
'petite verif
P_startH = P_startF + P_startS
If P_startH > 1 Then
    vd = MsgBox("OBLIGATOIR : p_startF + p_startS <= 1", 0, "GAFFE !")
    Exit Sub
End If
TAM(9).Text = Format(1 - P_startH) 'P_startVS

to_startF = Val(TAM(10).Text)
t_preS = Val(TAM(11).Text)
to1_startS = Val(TAM(12).Text)
t_preVS = Val(TAM(13).Text)  '70-150 ms
to1_startVS = Val(TAM(14).Text) '100 ms

t_pre_SlowDE = Val(TAM(15).Text)  't>10 ms ?
to_SlowDE = Val(TAM(16).Text)     '= to_startS = 30 ms ?

calcul_qoK (KqnZ1) '1 / 8.01) * Log(2800000 / Kqn) '1.9
TAM(17).Text = Format(qOK, "#.###")

to_Relax = Val(TAM(18).Text)  '0.5 'ms arret viscosite' pet etre fonction de de ht_step ?
t_pre_FastDE = Val(TAM(19).Text)
to_FastDE = Val(TAM(20).Text)
'Stop 'revoir dXz1 un peu partout car avant ds table T1T2met c'était X_a1
dXz1 = Val(TAM(21).Text)
If dXz1 < L_WS - aX Then 'on modifie
   dXz1 = L_WS - aX
   T1T2met.Edit: T1T2met("dXz1") = dXz1: T1T2met.Update
   TAM(21).Text = dXz1
End If

  X1 = aX / 2      'X_up
  X2 = -X1         'X_T
  X3 = X1 - L_WS   'X_down
  Xmin = aX - L_WS '-dX_E <0
  chi_p1 = 1 / Abs(X3)
    If dXz1 < 1 Then Stop  'oubli ?
  If dXz1 < Abs(Xmin) Then Stop 'fais gaffe !

 If Esc_Vi.RecordCount > 0 Then dbM.Execute "DELETE * FROM ESC_visco"
'epaisseur points/traits
e_pt = 8
e_tt = 2
```

```
'En abscisse le temps
  vx = "temps (ms)": xn = 0: xt = 5: xx = 45 'ms
'En ordonnée pT=T/T0
  yn = 0: yx = 1: yt = 0.1 '%
 'recherche d'erreur
  B = 0 '0=rien 1=Debug.print localisé
  sm = 1 'correspond à zp  = n° Escalier
  N_sm = 1 "correspond à k  = n° STAiR

If Escw.RecordCount > 0 Then dbM.Execute "delete * from ESC_verif2018"
'table ou sont stockee les tensions-STA de reference

' Fresc.Tag= "les 9", "V1", "V2", "V3", "1.5 nm", "3.4 nm", "4.3 nm")
'               0    1      2     3      4         5         6
If Fresc.Tag = 0 Then 'trace les 9
   'txV = "to_SB"
   'txV = "t_preSB"
    For zp = 1 To 9
        If txV = "t_preSB" Then
          t_preSB = 0
          'eassi 1
          'to1_SB = 1.5 'Choose(zp, 1.5, 1.5, 1.5, 1.5, 1.5, 1.5, 1.5, 1.5, 1.5) 'ref
          'to2_SB = Choose(zp, 4, 4, 4, 5, 5, 5, 5.9, 5.9, 5.9) 'ref
         'essai 2
          'to1_SB = Choose(zp, 1.2, 1.2, 1.2, 1.5, 1.5, 1.5, 1.5, 1.5, 1.5) 'ref
          'to2_SB = Choose(zp, 4.8, 4.8, 4.8, 5, 5, 5, 5.8, 5.8, 5.8) 'ref
        'eassi 3
          to1_SB = 1.5 'Choose(zp, 1.5, 1.5, 1.5, 1.5, 1.5, 1.5, 1.5, 1.5, 1.5) 'ref
          to2_SB = Choose(zp, 4.2, 4.2, 4.2, 5, 5, 5, 5.5, 5.5, 5.5) 'ref


        Else 'txV = "to_SB"
         Stop 'abandonné
          t_preSB = to1_df
          'Select Case zp
          '  Case Is <= 3: to1_SB = to1 ' + 6.3
          '  Case Is <= 6: to1_SB = to1 + 1.5   '1.2 '1.6
          '  Case Else: to1_SB = to1 + 2 '1.7 ' '2.2
          'End Select
          to1_SB = to1 + Choose(zp, 0, 1, 1, 2.5, 2.5, 2.5, 3, 3, 3)
        End If
        Esca.Seek "=", zp, -99: If Esca.NoMatch Then Stop
        vy = "-" + Format(Esca("ht_step")) + " nm," + Format(Esca("du_step")) + " ms"
        impscale X0 + Choose(zp, 0, 0, 0, -1, -1, -1, -2, -2, -2) * larg, Y0 + Choose(zp, 1, 2, 3, 1, 2, 3, 1, 2, 3) * haut + hentete, X0 + Choose(zp, 3,
3, 3, 2, 2, 2, 1, 1, 1) * larg, Y0 + Choose(zp, -2, -1, 0, -2, -1, 0, -2, -1, 0) * haut - hpage

        AAPH_ESC_trace_Ti_T1_pts 1
        AAPH_ESC_MON_calcul
        AAPH_ESC_MON_tracé
     Next 'zp

 Else  '
          For TRUC = 1 To 3 ' 3 cinétiques

           ' Fresc.Tag=  "V1", "V2", "V3", "1.5 nm", "3.4 nm", "4.3 nm")
           '             1      2     3      4         5         6

            Select Case TRUC
             Case 1 '
               zp = Choose(Fresc.Tag, 1, 2, 3, 1, 4, 7)
               Esca.Seek "=", zp, -99: If Esca.NoMatch Then Stop
               vy = "-" + Format(Esca("ht_step")) + " nm," + Format(Esca("du_step")) + " ms"
               impscale X0, Y0 + haut + hentete, X0 + larg, Y0 - 2 * haut - hpage

            Case 2
               zp = Choose(Fresc.Tag, 4, 5, 6, 2, 5, 8)
               Esca.Seek "=", zp, -99: If Esca.NoMatch Then Stop
               vy = "-" + Format(Esca("ht_step")) + " nm," + Format(Esca("du_step")) + " ms"
               impscale X0, Y0 + 2 * haut + hentete, X0 + larg, Y0 - haut - hpage

            Case 3
```

```
            zp = Choose(Fresc.Tag, 7, 8, 9, 3, 6, 9)
            Esca.Seek "=", zp, -99: If Esca.NoMatch Then Stop
            vy = "-" + Format(Esca("ht_step")) + " nm," + Format(Esca("du_step")) + " ms"
            impscale X0, Y0 + 3 * haut + hentete, X0 + larg, Y0 - hpage
        End Select

        AAPH_ESC_trace_Ti_T1_pts 1
        AAPH_ESC_MON_calcul
        AAPH_ESC_MON_tracé

    Next 'truc

 End If
End Sub
```

#### Sub AAPH_ESC_trace_Ti_T1_pts(iX As Integer)

```
'En abscisse le temps
'  vx = "temps (ms)": xn = 0: xt = 5: xx = 45 'ms
'En ordonnée pT=T/T0
'  yn = 0: yx = 1: yt = 0.1 '%

vd = ""
'vd = "echelle_2"
impenvi
With O
    Esca.Seek "=", zp, -99: If Esca.NoMatch Then Stop
    du_STEP_C = Esca("du_step") 'C pour Consigne / R pour Réel
    ht_step = -Esca("ht_step")
    .CurrentX = 15: .CurrentY = 116
    .FontSize = 11: .FontBold = -1
     'If txV = "t_preSB" Then
         tx4 = "to1_SB = " + Format(to1_SB) + "ms * to2_SB = " + Format(to2_SB)
     'Else
     '  tx4 = "to1_SB = " + Format(to1_SB)
     '  If to2_SB = 0 Then
     '    tx4 = tx4 + "ms * t_preSB = " + Format(t_preSB)
     '  Else
     '     tx4 = tx4 + "ms * to1_SB = " + Format(to1_SB)
     '  End If
     'End If
     O.Print "V = " + Format(ht_step / du_STEP_C, "0.##") + " nm/ms  *  " + tx4 + "ms"


If iX = 1 Then 'on trace les points

  'T0 etles marches d'escalier
    N_STEP = -1
    Esca.MoveNext
    Do While Esca("cas") = zp
        .DrawWidth = 5 * IIf(mmm = 1, 1, 4)
       'pas tracé pTi 'pour l'instant on travaille sur les %
       If IsNull(Esca("Ti_mm")) = 0 Then
          xav = du_STEP_C * Esca("pt"): xav = (xav - xn) * 100 / (xx - xn)
          yav = Esca("pTi") / 100: yav = (yav - yn) * 100 / (yx - yn)
          'O.PSet (xav, yav), QBColor(12) 'coul
       End If
       'pas tracé pT1
       If IsNull(Esca("T1_mm")) = 0 Then
          xmi = du_STEP_C * (Esca("pt") - 1): xmi = (xmi - xn) * 100 / (xx - xn)
          ymi = Esca("pT1") / 100: ymi = (ymi - yn) * 100 / (yx - yn)
          'O.PSet (xmi, ymi), QBColor(1) 'coul
       End If

       'exponentielles en serie
       If Esca("pt") > 0 Then
          N_STEP = N_STEP + 1

        'd'abord les points
        xav = Esca("tps_0"): xav = (xav - xn) * 100 / (xx - xn)
        yav = Esca("pTin_0") / 100: yav = (yav - yn) * 100 / (yx - yn)
        .DrawWidth = 8 * IIf(mmm = 1, 1, 4)
        O.PSet (xav, yav), QBColor(12) 'coul
        .CurrentX = xav
        .CurrentY = yav
        For z = 1 To 11
            vx = Format(z)
            If IsNull(Esca("Tin_" + vx + "_mm")) = 0 Then
                  xav = Esca("tps_" + vx): xav = (xav - xn) * 100 / (xx - xn)
                  yav = Esca("pTin_" + vx) / 100: yav = (yav - yn) * 100 / (yx - yn)
                  .DrawWidth = 4 * IIf(mmm = 1, 1, 4)
                  O.PSet (xav, yav), QBColor(1) 'coul
            End If
         Next 'z

        'ensuite les droites
         TE = ""
```

```
        If TE = "pts_relies" Then
           .DrawWidth = 1 * IIf(mmm = 1, 1, 4)
            xav = Esca("tps_0"): xav = (xav - xn) * 100 / (xx - xn)
            yav = Esca("pTin_0") / 100 ': yav = (yav - yn) * 100 / (yx - yn)
            .DrawWidth = 1 * IIf(mmm = 1, 1, 4)
            .CurrentX = xav
            .CurrentY = yav
            For z = 1 To 11
                vx = Format(z)
                If IsNull(Esca("Tin_" + vx + "_mm")) = 0 Then
                      xav = Esca("tps_" + vx): xav = (xav - xn) * 100 / (xx - xn)
                      yav = Esca("pTin_" + vx) / 100: yav = (yav - yn) * 100 / (yx - yn)
                      O.Line -(xav, yav), QBColor(0)
                End If
             Next 'z
       End If
      End If


      Esca.MoveNext: If Esca.EOF Then Exit Do
   Loop
   If vd = "echelle_2" Then  'ligne horizontale sur dernier point
      .DrawWidth = IIf(mmm = 1, 1, 5)
       O.Line (0, yav)-(100, yav)
   End If

ElseIf iX = 2 Then 'on trace q pour le suivi de la viscosité
   'O.Line (0, 0)-(100, 0)

   .DrawWidth = 7 * IIf(mmm = 1, 1, 4)
   Esc_Vi.Seek "=", zp, 1, -1: If Esc_Vi.NoMatch Then Stop
   ht_step = Esc_Vi("dX_stairT")
   du_STEP_C = Esc_Vi("to_stairT")
   Do While Esc_Vi("cas") = zp


      'xav = (Esc_Vi("t_finstair") / du_STEP_C) * ht_step
      xav = Esc_Vi("t_finstair")
      xav = (xav - xn) * 100 / (xx - xn)

     'calcul avec pente
      yav = Esc_Vi(IIf(vx = "q", "q_pente", "Kqn"))
      yav = (yav - yn) * 100 / (yx - yn)
      .DrawWidth = 6 * IIf(mmm = 1, 1, 4)
      O.PSet (xav, yav), QBColor(0) 'coul

     'calcul avec q=q_T0+log(pTi)/log(nhs)
      yav = Esc_Vi(IIf(vx = "q", "q_log", "K_log"))
      yav = (yav - yn) * 100 / (yx - yn)

      'O.PSet (xav, yav), QBColor(13) 'coul
       .DrawWidth = IIf(mmm = 1, 1, 4)
      O.Line (xav - 1, yav)-(xav + 1, yav), QBColor(13)
      O.Line (xav, yav - 1)-(xav, yav + 1), QBColor(13)
      Esc_Vi.MoveNext
      If Esc_Vi.EOF Then Exit Do

   Loop

   '.DrawWidth = IIf(mmm = 1, 1, 4)
   ' yav = IIf(vx = "q", 2, 1.1)
   ' yav = (yav - yn) * 100 / (yx - yn)
   ' O.Line (0, yav)-(100, yav), QBColor(12)
   yav = 10
   O.Line (0, yav)-(100, yav)
End If
'Stop
'vd = ""
'vd = "echelle_2"
If vd = "echelle_2" Then '2eme echelle de nm
   'on trace sur l'ordonné yav une nouvelle echelle
```

```
    'npas = Int((xx - xn) / xt)
    'titre
    tx1 = "dist(nm)"
    .FontBold = -1: .FontSize = IIf(mmm = 1, 8, 7)
    .CurrentX = 102 - .TextWidth(tx1) '91

   Select Case iX
    Case 1:  'pts Ti
      .CurrentY = yav + Choose(zp, -7, -7, -7, 8, 14, 14, 8, 14, 14)
     Case 2 'visco
      .CurrentY = yav - 7
    End Select
    O.Print tx1

    .FontSize = IIf(mmm = 1, 8, 7)
    .FontBold = 0
    npas = Int((xx - xn) / xt + 0.5)

    For z = Int(xn / xt) To npas + Int(xn / xt)
        xav = (z * xt - xn) * 100 / (xx - xn)

        Select Case iX
         Case 1:  'pts Ti
          O.Line (xav, yav)-(xav, yav + Choose(zp, -2, -2, -2 - 2, -2, 2, 2, -2, 2, 2))
          .CurrentY = yav + 1.5 * Choose(zp, -1, -1, -1 - 1, -1, 5, 5, -1, 5, 5)
        Case 2 'visco
            O.Line (xav, yav)-(xav, yav - 2)
          .CurrentY = yav - 1.5
        End Select

        .CurrentX = xav - 0.7
        P = z * (xt / du_STEP_C) * ht_step
        O.Print Format(P, "#0.#")
    Next 'z
End If

'If Ix = 1 Then   'on trace L_WS
    coul = QBColor(IIf(mmm = 1, 7, 0))
    .DrawWidth = 2 * IIf(mmm = 1, 1, 5)
    'xav = (L_WS * du_STEP_C / ht_step - xn) * 100 / (xx - xn)
    'O.Line (xav, 0)-(xav, 100), coul
    'If drap = 1 Then .CurrentX = xav: .CurrentY = 100: O.Print "L_WS"


    xap = L_WS + L_preWS
    xav = (xap * du_STEP_C / ht_step - xn) * 100 / (xx - xn)
    O.Line (xav, 0)-(xav, 100), coul
    .CurrentX = xav: .CurrentY = 100: O.Print Format(xap) + "nm"
    ' on trace 20nm
    '  xav = (20 * du_STEP_C / ht_step - xn) * 100 / (xx - xn)
    'O.Line (xav, 0)-(xav, 100), coul
'End If

End With
End Sub
```

#### Sub AAPH_ESC_MON_calcul() 'octobre

```
'****************** RECUP  VAL issues de table Esca(les points)  et remplissage table escw (durée et pTi)
Esca.Seek "=", zp, -99 'zp = n° d'ESCalier ou de graphiques de 1 à 9

ht_step = Abs(Esca("ht_step"))
du_STEP_C = Esca("du_step")

N_STAiR_ds_dX_Max = Int(L_WS / ht_step)
Morceau_STAiR_ds_dX_Max = L_WS / ht_step - N_STAiR_ds_dX_Max

N_STAiR_ds_dX_T = Int(dX_T / ht_step)
Morceau_STAiR_ds_dX_T = dX_T / ht_step - N_STAiR_ds_dX_T

'comme on ne sait pas trop ce qui se passe on arrondit à l'entier supeirut
N_STAiR_ds_dX_Pre = Int(L_preWS / ht_step) 'N_etude = N_STAiR_ds_dX_Pre
Morceau_STAiR_ds_dX_Pre = L_preWS / ht_step - N_STAiR_ds_dX_Pre  'Morceau_STAiR_ds_dX_Pre

'N_clik = N_STAiR_ds_dX_Pre + N_STAiR_ds_dX_Max + IIf(Morceau_STAiR_ds_dX_Max = 0, 0, 1)
N_clik = N_STAiR_ds_dX_Max + IIf(Morceau_STAiR_ds_dX_Max = 0, 0, 1)

' du_STAiR_V(1 To 15) As Single 'V por Vrai
For k = 1 To 15: du_STAiR_V(k) = 0: Next

k = 1 'on initialise le compteur de marches (stair)
t_debSTEP = 0 'por recuperer t_finSTEP

Esca.Seek "=", zp, 1: If Esca.NoMatch Then Stop
Do While Esca("cas") = zp 'on deroule pour chaque STEP
    For z = 1 To 11
        If IsNull(Esca("pTin_" + Format(z))) Then Exit For
    Next 'z
    If z < 3 Then 'fin de l'escalier: on compte de nre de marches et on sort
      N_STEP = k - 1
      Exit Do
    End If

    t_finSTEP = Esca("tps_" + Format(z - 1))
    t_fin_Phase1 = Esca("tps_0")

    Escw.AddNew
     Escw("cas") = Esca("cas")
     Escw("esc") = zp
     Escw("stair") = k
     Escw("VAR_typ") = -1
     Escw("VAR_nom") = "DATA ori"
     Escw("dX_stairT") = ht_step
     Escw("to_stairT") = du_STEP_C
     Escw("to_stairM") = t_finSTEP - t_debSTEP
     Escw("dX_ESC") = ht_step * k
     Escw("t_ESC") = du_STEP_C * k
     Escw("t_P1") = t_fin_Phase1 - t_debSTEP
     Escw("t_finP1") = t_fin_Phase1
     Escw("t_debSTAiR") = t_debSTEP
     Escw("t_finSTAiR") = t_finSTEP
     du_STAiR_V(k) = t_finSTEP - t_fin_Phase1 'la vriar duée pendat laquelle les remontees se font
     Escw("Tot") = Null
     For i = 1 To 30:  Escw("d" + Format(i)) = Null: Next
    Escw.Update

    'step suivant
    Esca.MoveNext
    k = k + 1 'compteur de marches
    t_debSTEP = t_finSTEP

 Loop
****************************************************************************************************** WS
ISOMETRIE
' 1 *************************** tete en WS en isometrie
'ATENTION on commence ne isometrie k=0
For k = 0 To N_STEP  'nbre de marches experimentales
```

```
    Escw.AddNew
     Escw("cas") = Esca("cas")
     Escw("esc") = zp
     Escw("stair") = k  ' en fait marche precedente
     Escw("VAR_typ") = 0
     Escw("VAR_nom") = "WS Isom"
     Escw("dX_stairT") = ht_step
     Escw("to_stairT") = du_STEP_C
     Escw("to_stairM") = Null
     Escw("dX_ESC") = ht_step * k
     Escw("t_ESC") = du_STEP_C * k
     Escw("t_P1") = Null: Escw("t_finP1") = Null: Escw("t_debSTAiR") = Null: Escw("t_finSTAiR") = Null
     For i = 1 To 30:  Escw("d" + Format(i)) = Null: Next

     sY = 0
     If (dX_T + k * ht_step) <= L_WS Then ' tout les TEtes en WS_tetanos encore presentes
        drap = 0
        sN = k + N_STAiR_ds_dX_T
     ElseIf k * ht_step <= L_WS Then
         drap = 1
         sN = N_STAiR_ds_dX_Max
     Else
        drap = -1
     End If

     If drap > -1 Then
        For i = (k + 1) To sN
           Xa = X1 - (i - 1) * ht_step
           Xb = X1 - i * ht_step
           pT = (ht_step / dX_T) * (1 + 0.5 * (Xa + Xb) / Abs(X3))
           Escw("d" + Format(i)) = pT
           sY = sY + pT
        Next 'i
        If drap = 0 And Morceau_STAiR_ds_dX_T > 0 Then 'le morceau est devant  donc i=sN+1
           'i = sN + 1
           Xa = X1 - (i - 1) * ht_step
           Xb = Xa - Morceau_STAiR_ds_dX_T * ht_step
           If Xb < X3 Then Xb = X3 ' Stop 'pas normal car drap=0 (voir + haut)
           pT = (Morceau_STAiR_ds_dX_T * ht_step / dX_T) * (1 + 0.5 * (Xa + Xb) / Abs(X3))
           Escw("d" + Format(i)) = pT
           sY = sY + pT
        ElseIf drap = 1 And Morceau_STAiR_ds_dX_Max > 0 Then
           'i= sN+1 = N_STAiR_ds_dX_Max +1
           Xa = X1 - (i - 1) * ht_step
           If Xa < X3 Then Stop
           Xb = X3
           If Abs((Xa - Xb) - Morceau_STAiR_ds_dX_Max * ht_step) > 0.000001 Then Stop
           pT = (Morceau_STAiR_ds_dX_Max * ht_step / dX_T) * (1 + 0.5 * (Xa + Xb) / Abs(X3))
           Escw("d" + Format(i)) = pT
           sY = sY + pT
       End If
     End If
     Escw("Tot") = sY
   Escw.Update
Next 'k
```

```
' ************************************************************LA *****************************************************
REMONTEES
'je ne travaille que les stair_ds_dXpre_entieres : le morceau qui suit est traité avec les NEW
' 11 12 ----------------------remontee rapide tete préalablment en SB (t=0) en isometrie
For F = 1 To 2 '1=Fast 2=Slow
 For k = 1 To N_STEP 'nbre de marches experimentales
    Escw.AddNew
     Escw("cas") = Esca("cas")
     Escw("esc") = zp
     Escw("stair") = k
     Escw("VAR_typ") = 10 + F
     Escw("VAR_nom") = "WS+Rem" + IIf(F = 1, "F", "S") + " t=0"
     Escw("dX_stairT") = ht_step
     Escw("to_stairT") = du_STEP_C
     Escw("to_stairM") = Null
     Escw("dX_ESC") = ht_step * k
     Escw("t_ESC") = du_STEP_C * k
     Escw("t_P1") = Null: Escw("t_finP1") = Null: Escw("t_debSTAiR") = Null: Escw("t_finSTAiR") = Null
     For i = 1 To 30:  Escw("d" + Format(i)) = Null: Next

     drap = 0
     If k <= N_STAiR_ds_dX_Pre Then 'N_etude = N_STAiR_ds_dX_Pre + 0_ou_1
        N_F = 1
        sN = k
     ElseIf k <= N_STAiR_ds_dX_Max Then
        N_F = k - N_STAiR_ds_dX_Pre + 1
        sN = k
        'drap = 1 'Morceau_ds_dX_pre
     ElseIf k <= N_STAiR_ds_dX_Max + N_STAiR_ds_dX_Pre Then
        N_F = k - N_STAiR_ds_dX_Pre + 1
        sN = N_STAiR_ds_dX_Max
        drap = 2 'Morceau_ds_dX_Max
     Else
        drap = -1
     End If

     sY = 0
     If drap > -1 Then AAPH_ESC_MON_calcul_pT_WS_Rem_ISO_t_0 1
     Escw("Tot") = sY

   Escw.Update
 Next 'k
Next 'F
'Stop

' 13 14 ----------------------remontee rapide et lente tete NEW

For F = 1 To 2 '1=Fast 2=Slow
 For k = 1 To N_STEP 'nbre de marches experimentales
    Escw.AddNew
     Escw("cas") = Esca("cas")
     Escw("esc") = zp
     Escw("stair") = k
     Escw("VAR_typ") = 12 + F
     Escw("VAR_nom") = "WS+Rem" + IIf(F = 1, "F", "S") + " NEW"
     Escw("dX_stairT") = ht_step
     Escw("to_stairT") = du_STEP_C
     Escw("to_stairM") = Null
     Escw("dX_ESC") = ht_step * k
     Escw("t_ESC") = du_STEP_C * k
     Escw("t_P1") = Null: Escw("t_finP1") = Null: Escw("t_debSTAiR") = Null: Escw("t_finSTAiR") = Null
     For i = 1 To 30:  Escw("d" + Format(i)) = Null: Next
```

```
    drap = 0
    If k <= N_STAiR_ds_dX_Pre Then 'rien
      drap = -1
    ElseIf k <= N_STAiR_ds_dX_Max + N_STAiR_ds_dX_Pre Then
      N_F = 1
      sN = k - N_STAiR_ds_dX_Pre
      drap = 1 'la stair N° sN contient le Morceau_ds_dXpre en SB à t=0 avec p_statF
   Else
      N_F = 1
      sN = N_STAiR_ds_dX_Max
      drap = 2 ' on est en regime répétifit
    End If
    sY = 0
    If drap > -1 Then AAPH_ESC_MON_calcul_pT_WS_rem_NEW 1
    Escw("Tot") = sY
  Escw.Update
 Next 'k
Next 'F
'Stop


' 15 ------TOUUUUUT -----------------remontee tout
zB = 15
For k = 1 To N_STEP 'nbre de marches experimentales
   Escw.AddNew
    Escw("cas") = Esca("cas")
    Escw("esc") = zp
    Escw("stair") = k
    Escw("VAR_typ") = zB
    Escw("VAR_nom") = "WS+Rem TOUT"
    Escw("dX_stairT") = ht_step
    Escw("to_stairT") = du_STEP_C
    Escw("to_stairM") = Null
    Escw("dX_ESC") = ht_step * k
    Escw("t_ESC") = du_STEP_C * k
    Escw("t_P1") = Null: Escw("t_finP1") = Null: Escw("t_debSTAiR") = Null: Escw("t_finSTAiR") = Null
    For i = 1 To 30:  Escw("d" + Format(i)) = Null: Next
    Escw.Update

  sY = 0
  For F = 1 To 2 '1=Fast 2=Slow
      Escw.Seek "=", zp, k, zB: If Escw.NoMatch Then Stop
      Escw.Edit

     'teten en SB à t=0
      drap = 0
      If k <= N_STAiR_ds_dX_Pre Then
        N_F = 1
        sN = k
      ElseIf k <= N_STAiR_ds_dX_Max Then
        N_F = k - N_STAiR_ds_dX_Pre + 1
        sN = k
      ElseIf k <= N_STAiR_ds_dX_Max + N_STAiR_ds_dX_Pre Then
        N_F = k - N_STAiR_ds_dX_Pre + 1
        sN = N_STAiR_ds_dX_Max
        drap = 2 'Morceau_ds_dX_Max
      Else
        drap = -1
      End If

      If drap > -1 Then AAPH_ESC_MON_calcul_pT_WS_Rem_ISO_t_0 3' on somme
```

```
      'teten NEW
        drap = 0
        If k <= N_STAiR_ds_dX_Pre Then 'rien
            drap = -1
        ElseIf k <= N_STAiR_ds_dX_Max + N_STAiR_ds_dX_Pre Then
            N_F = 1
            sN = k - N_STAiR_ds_dX_Pre
            drap = 1 'la stair N° sN contient le Morceau_ds_dXpre en SB à t=0 avec p_statF
        Else
            N_F = 1
            sN = N_STAiR_ds_dX_Max
            drap = 2 ' on est en regime répétifit
        End If
        If drap > -1 Then AAPH_ESC_MON_calcul_pT_WS_rem_NEW 3 'on somme

        Escw.Update
    Next 'F

    Escw.Seek "=", zp, k, zB: If Escw.NoMatch Then Stop
    Escw.Edit: Escw("Tot") = sY:  Escw.Update
Next 'k
'Stop


'**********************ICI*****************************pT1 elas (valeurs pt à la fin de la phase 1 de la marche
' 1 2 ----------------------  pT1 elas WS_Fast and slow à t=en SB à t=0
For F = 1 To 2 '1=Fast 2=Slow
 For k = 1 To N_STEP 'nbre de marches experimentales
    Escw.AddNew
     Escw("cas") = Esca("cas")
     Escw("esc") = zp
     Escw("stair") = k
     Escw("VAR_typ") = F
     Escw("VAR_nom") = "WS_" + IIf(F = 1, "F", "S") + " t=0"
     Escw("dX_stairT") = ht_step
     Escw("to_stairT") = du_STEP_C
     Escw("to_stairM") = Null
     Escw("dX_ESC") = ht_step * k
     Escw("t_ESC") = du_STEP_C * k
     Escw("t_P1") = Null: Escw("t_finP1") = Null: Escw("t_debSTAiR") = Null: Escw("t_finSTAiR") = Null
     For i = 1 To 30:  Escw("d" + Format(i)) = Null: Next

     drap = 0
     If k = 1 Then 'step 1 : rien car n'est concerne que les WS de l'isometrie du départ
        drap = -1
     ElseIf k <= N_STAiR_ds_dX_Pre + 1 Then ''on decale tout de 1 et le Morceau qui reste n'est pas incriminé0
        N_F = 2 '1
        sN = k
     ElseIf k <= N_STAiR_ds_dX_Max Then 'il faut tenir du bout qui reste : drpa= 1
        N_F = k - N_STAiR_ds_dX_Pre + 1
        sN = k
     ElseIf k <= N_STAiR_ds_dX_Max + N_STAiR_ds_dX_Pre Then
        N_F = k - N_STAiR_ds_dX_Pre + 1
        sN = N_STAiR_ds_dX_Max
        drap = 2
     Else
        drap = -1
     End If
     sY = 0

     If drap > -1 Then AAPH_ESC_MON_calcul_pT1_elas_ISO_t_0 1
     Escw("Tot") = sY

   Escw.Update
 Next 'k
Next 'F
'Stop
' 3 4 ----------------------pT1 elas WS_Fast and slow  NEW
For F = 1 To 2 '1=Fast 2=Slow

 For k = 1 To N_STEP 'nbre de marches experimentales
```

```
   Escw.AddNew
    Escw("cas") = Esca("cas")
    Escw("esc") = zp
    Escw("stair") = k
    Escw("VAR_typ") = 2 + F
    Escw("VAR_nom") = "WS_ " + IIf(F = 1, "F", "S") + " NEW"
    Escw("dX_stairT") = ht_step
    Escw("to_stairT") = du_STEP_C
    Escw("to_stairM") = Null
    Escw("dX_ESC") = ht_step * k
    Escw("t_ESC") = du_STEP_C * k
    Escw("t_P1") = Null: Escw("t_finP1") = Null: Escw("t_debSTAiR") = Null: Escw("t_finSTAiR") = Null
    For i = 1 To 30:  Escw("d" + Format(i)) = Null: Next

    drap = 0
    If k <= N_STAiR_ds_dX_Pre + 1 Then
       drap = -1
    ElseIf k <= N_STAiR_ds_dX_Max + N_STAiR_ds_dX_Pre Then
       N_F = 2
       sN = k - N_STAiR_ds_dX_Pre
       drap = 1 'la stair N° sN contient le Morceau_ds_dXpre en SB à t=0 avec p_statF
    Else
       N_F = 2
       sN = N_STAiR_ds_dX_Max
       drap = 2 ' on est en regime répétifit
    End If

    sY = 0
    If drap > -1 Then AAPH_ESC_MON_calcul_pT1_elas_NEW 1
    Escw("Tot") = sY

  Escw.Update
 Next 'k
Next 'F

'Stop
```

```
' 5 ------TOUUUUUT -----------------pT1 elas WS_TOUT
zB = 5
For k = 1 To N_STEP 'nbre de marches experimentales
    Escw.AddNew
     Escw("cas") = Esca("cas")
     Escw("esc") = zp
     Escw("stair") = k
     Escw("VAR_typ") = zB
     Escw("VAR_nom") = "WS_F_S TOUT"
     Escw("dX_stairT") = ht_step
     Escw("to_stairT") = du_STEP_C
     Escw("to_stairM") = Null
     Escw("dX_ESC") = ht_step * k
     Escw("t_ESC") = du_STEP_C * k
     Escw("t_P1") = Null: Escw("t_finP1") = Null: Escw("t_debSTAiR") = Null: Escw("t_finSTAiR") = Null
     For i = 1 To 30:  Escw("d" + Format(i)) = Null: Next
     Escw.Update

  sY = 0
  For F = 1 To 2 '1=Fast 2=Slow
        Escw.Seek "=", zp, k, zB: If Escw.NoMatch Then Stop
        Escw.Edit
      'teten en SB à t=0
       drap = 0
        If k = 1 Then 'step 1 : rien car n'est concerne que les WS de l'isometrie du départ
           drap = -1
        ElseIf k <= N_STAiR_ds_dX_Pre + 1 Then "on decale tout de 1 et le Morceau qui reste n'est pas incriminé0
           N_F = 2 '1
           sN = k
        ElseIf k <= N_STAiR_ds_dX_Max Then 'il faut tenir du bout qui reste : drpa= 1
           N_F = k - N_STAiR_ds_dX_Pre + 1
           sN = k
        ElseIf k <= N_STAiR_ds_dX_Max + N_STAiR_ds_dX_Pre Then
           N_F = k - N_STAiR_ds_dX_Pre + 1
           sN = N_STAiR_ds_dX_Max
           drap = 2
        Else
           drap = -1
        End If

        '****************************
        If drap > -1 Then AAPH_ESC_MON_calcul_pT1_elas_ISO_t_0 3'on somme

      'teten NEW
         drap = 0
         If k <= N_STAiR_ds_dX_Pre + 1 Then
             drap = -1
         ElseIf k <= N_STAiR_ds_dX_Max + N_STAiR_ds_dX_Pre Then
             N_F = 2
               sN = k - N_STAiR_ds_dX_Pre
               drap = 1 'la stair N° sN contient le Morceau_ds_dXpre en SB à t=0 avec p_statF
         Else
               N_F = 2
               sN = N_STAiR_ds_dX_Max
               drap = 2 ' on est en regime répétifit
         End If

        '**************************
        If drap > -1 Then AAPH_ESC_MON_calcul_pT1_elas_NEW 3'on somme
        Escw.Update
    Next 'F

    Escw.Seek "=", zp, k, zB: If Escw.NoMatch Then Stop
    Escw.Edit: Escw("Tot") = sY:  Escw.Update
Next 'k

'Stop
' ******************************************************************************************* FastDE
'*********  - pT du aux detachment rapide a faire quant role viscosité appréhenée

' ******************************************************************************************* pT1
```

```
' *************** pT1 elas
zB = 32
For k = 1 To N_STEP 'nbre de marches experimentales
    Escw.AddNew
     Escw("cas") = Esca("cas")
     Escw("esc") = zp
     Escw("stair") = k
     Escw("VAR_typ") = zB
     Escw("VAR_nom") = "pT1 elas"
     Escw("dX_stairT") = ht_step
     Escw("to_stairT") = du_STEP_C
     Escw("to_stairM") = Null
     Escw("dX_ESC") = ht_step * k
     Escw("t_ESC") = du_STEP_C * k
     Escw("t_P1") = Null: Escw("t_finP1") = Null: Escw("t_debSTAiR") = Null: Escw("t_finSTAiR") = Null
     For i = 1 To 30:  Escw("d" + Format(i)) = Null: Next
    Escw.Update

     Escw.Seek "=", zp, k, 0: If Escw.NoMatch Then Stop
     sY = Escw("Tot")
     Escw.Seek "=", zp, k, 5: If Escw.NoMatch Then Stop
     sY = sY + Escw("tot")

   Escw.Seek "=", zp, k, zB: If Escw.NoMatch Then Stop
   Escw.Edit: Escw("Tot") = sY: Escw.Update
Next 'k

' *************** pTi
zB = 33
For k = 1 To N_STEP 'nbre de marches experimentales
    Escw.AddNew
     Escw("cas") = Esca("cas")
     Escw("esc") = zp
     Escw("stair") = k
     Escw("VAR_typ") = zB
     Escw("VAR_nom") = "pTi"
     Escw("dX_stairT") = ht_step
     Escw("to_stairT") = du_STEP_C
     Escw("to_stairM") = Null
     Escw("dX_ESC") = ht_step * k
     Escw("t_ESC") = du_STEP_C * k
     Escw("t_P1") = Null: Escw("t_finP1") = Null: Escw("t_debSTAiR") = Null: Escw("t_finSTAiR") = Null
     For i = 1 To 30:  Escw("d" + Format(i)) = Null: Next
    Escw.Update

     Escw.Seek "=", zp, k, 0: If Escw.NoMatch Then Stop 'WS isom
     sY = Escw("Tot")
     Escw.Seek "=", zp, k, 15: If Escw.NoMatch Then Stop 'WS+rem tout
     sY = sY + Escw("tot")

   Esca.Seek "=", zp, k: If Esca.NoMatch Then Stop
   courbe = Choose(zp, 6, 5, 5, 8, 7, 6, 8, 8, 7) 'rentré à ma main LA barbe !
   sYY = Esca("pTin_" + Format(courbe)) / 100

   Escw.Seek "=", zp, k, zB: If Escw.NoMatch Then Stop
   Escw.Edit
     Escw("Tot") = sY 'pTi calculé
     Escw("d1") = sYY 'pTi mesuré
   Escw.Update
Next 'k
'*********  pT1 VISCO+Elas mesuré = Esca("pTin_0")
zB = 31
For k = 1 To N_STEP 'nbre de marches experimentales
     Escw.AddNew
     Escw("cas") = Esca("cas")
     Escw("esc") = zp
     Escw("stair") = k
     Escw("VAR_typ") = zB
     Escw("VAR_nom") = "pT1 visco+elas"
     Escw("dX_stairT") = ht_step
```

```
    Escw("to_stairT") = du_STEP_C
    Escw("to_stairM") = Null
    Escw("dX_ESC") = ht_step * k
    Escw("t_ESC") = du_STEP_C * k
    Escw("t_P1") = Null: Escw("t_finP1") = Null: Escw("t_debSTAiR") = Null: Escw("t_finSTAiR") = Null: Escw("Tot") = Null
    For i = 1 To 30:  Escw("d" + Format(i)) = Null: Next
  Escw.Update

  'pT1 mesuré
   Esca.Seek "=", zp, k: If Esca.NoMatch Then Stop
   pT_pas = Esca("pTin_0") / 100

  'pT1 theorique
    If k = 1 Then
       pTi = 1
       Escw.Seek "=", zp, k, 32: If Escw.NoMatch Then Stop  'WS isom

      ht_step = Escw("dX_stairT")
      pT1_elas = Escw("Tot")
      pz1_ssV = (pTi - pT1_elas) / ht_step
      'If Abs(pz1_ssV - 0.133) > 0.001 Then Stop
      Select Case zp
       Case Is <= 3: KqnZ1 = 1.8
       Case Is <= 6: KqnZ1 = 1.64 '1.64 '1.62
       Case Else: KqnZ1 = 1.5 '1.5
      End Select
      calcul_qoK (KqnZ1)
      qZ1 = qOK

    Else
       Escw.Seek "=", zp, k - 1, 33: If Escw.NoMatch Then Stop 'WS isom
       pTi = Escw("Tot")
       qZ1 = qOK + Log(pTi) / Log(N_hs)
       KqnZ1 = calcul_Kqn(qZ1)
    End If
    'If k = 1 Then KqnZ1 = 1.6
    If zp <= 3 And k > 1 Then
       Kqn = Choose(zp, 2.9, 3.35, 3.85)
       z = N_clik + 1
       If k > z Then
          KqnZ1 = Kqn
       Else 'on effectue une descente lineaire entre 1.8 et kqn les données au dessus
          KqnZ1 = 1.8 - (k - 1) * (1.8 - Kqn) / (z - 1)
       End If
    End If
    pT1 = 1 - ht_step * KqnZ1 * pz1_ssV
    If pT1 < 0 Then pT1 = 0
  'on se repalce et on stocke
   Escw.Seek "=", zp, k, zB: If Escw.NoMatch Then Stop 'WS isom
    Escw.Edit
     Escw("d1") = pT_pas ' pT1 mesuré
     Escw("d3") = qZ1
     Escw("d4") = KqnZ1
     Escw("Tot") = pT1 'pT1 calcilé
    Escw.Update
 Next 'k
```

#### Sub AAPH_ESC_MON_calcul_pT_WS_Rem_ISO_t_0(Icod As Integer)

```
'WS+ Rem : Icod= 1  1 valeur   Icd= 3 on somme

'mise à 0 pour etre tranquille
For i = 1 To 15: Sdu_STAiR_V(i) = 0: Next
'faut inverser lstair 1 correspond à la durée k  etc
Sdu_STAiR_V(1) = du_STAiR_V(k)

If k <= N_clik Then
  For i = (k - 1) To 1 Step -1
     TRUC = k - i + 1
     Sdu_STAiR_V(TRUC) = Sdu_STAiR_V(TRUC - 1) + du_STAiR_V(i)

  Next 'i
```

```
Else 'faut décaler en fonction de n_clik
  For i = (k - 1) To (k - N_clik + 1) Step -1
     TRUC = k - i + 1
     Sdu_STAiR_V(TRUC) = Sdu_STAiR_V(TRUC - 1) + du_STAiR_V(i)
  Next 'i
End If

For i = N_F To sN
    Xa = X1 - (i - 1) * ht_step
    Xb = X1 - i * ht_ste

    If F = 1 Then 'Fast en SB à t= 0
         Qdp = P_startF * (1 - Exp(-Sdu_STAiR_V(i) / to_startF))
    Else 'slow en SB à t=0
        Qdp = P_startS * (1 - Exp(-Sdu_STAiR_V(i) / to1_startS))
    End If

      'sm = 1 'correspond à zp  = n° Escalier      'N_sm = 1 "correspond à k  = n° STAiR
     pT = Qdp * (ht_step / dX_T) * (1 + 0.5 * (Xa + Xb) / Abs(X3))

     'If F = 2 And i = 2 And k = 8 Then Stop

     vd = "d" + Format(i)
     If Icod = 1 Or (Icod = 3 And F = 1) Then
        Escw(vd) = pT
     Else
        Escw(vd) = pT + Escw(vd)
     End If
     sY = sY + pT
 Next 'i

 If drap = 2 And Morceau_STAiR_ds_dX_Max > 0 Then
        If i <> (N_STAiR_ds_dX_Max + 1) Then Stop
        Xa = X1 - (i - 1) * ht_step
        Xb = X3

        If F = 1 Then 'Fast en SB à t= 0
             Qdp = P_startF * (1 - Exp(-Sdu_STAiR_V(i) / to_startF))
        Else 'slow en SB à t=0
            Qdp = P_startS * (1 - Exp(-Sdu_STAiR_V(i) / to1_startS))
        End If

         pT = Qdp * (Morceau_STAiR_ds_dX_Max * ht_step / dX_T) * (1 + 0.5 * (Xa + Xb) / Abs(X3))
       vd = "d" + Format(i)
       If Icod = 1 Or (Icod = 3 And F = 1) Then
            Escw(vd) = pT
        Else
            Escw(vd) = pT + Escw(vd)
        End If
        sY = sY + pT
  End If

End Sub
```

```
Sub AAPH_ESC_MON_calcul_pT_WS_rem_NEW(Icod As Integer)
'WS+ Rem : Icod= 1  1 valeur   Icd= 3 on somme
If N_F > sN Then Stop '?

'mise à 0 pour etre tranquille
For i = 1 To 15: Sdu_STAiR_V(i) = 0: Next
'faut inverser lstair 1 correspond à la durée k  etc
Sdu_STAiR_V(1) = du_STAiR_V(k)

If k <= N_clik Then

  For i = (k - 1) To 1 Step -1
     TRUC = k - i + 1
     Sdu_STAiR_V(TRUC) = Sdu_STAiR_V(TRUC - 1) + du_STAiR_V(i)
  Next 'i

Else 'faut décaler en fonction de n_clik

  For i = (k - 1) To (k - N_clik + 1) Step -1
     TRUC = k - i + 1
     Sdu_STAiR_V(TRUC) = Sdu_STAiR_V(TRUC - 1) + du_STAiR_V(i)
  Next 'i
End If

'pre
Rds = 0
If to2_SB = 0 Then 'simple exponentielle
    Pdp = P_startF * (1 - Exp(-(Sdu_STAiR_V(N_STAiR_ds_dX_Pre) - t_preSB) / to1_SB))
   If Morceau_STAiR_ds_dX_Pre > 0 Then 'Morceau ds dX_pre
     Rds = P_startF * (1 - Exp(-(Sdu_STAiR_V(N_STAiR_ds_dX_Pre + 1) - t_preSB) / to1_SB))
   End If

Else '* ** double exponentizelles
   sX = to1_SB * Exp(-(Sdu_STAiR_V(N_STAiR_ds_dX_Pre) - t_preSB) / to1_SB) / (to1_SB - to2_SB)
   sXX = to2_SB * Exp(-(Sdu_STAiR_V(N_STAiR_ds_dX_Pre) - t_preSB) / to2_SB) / (to2_SB - to1_SB)
   Pdp = P_startF * (1 - sX - sXX)
   If Morceau_STAiR_ds_dX_Pre > 0 Then
     sX = to1_SB * Exp(-(Sdu_STAiR_V(N_STAiR_ds_dX_Pre + 1) - t_preSB) / to1_SB) / (to1_SB - to2_SB) 'Morceau ds dX_pre
     sXX = to2_SB * Exp(-(Sdu_STAiR_V(N_STAiR_ds_dX_Pre + 1) - t_preSB) / to2_SB) / (to2_SB - to1_SB) 'Morceau ds dX_pre
     Rds = P_startF * (1 - sX - sXX)
   End If
End If

For i = N_F To sN
     ' STAiR = [Xb Xc]_RDS + (Xc ; Xa]_PdP
     Xa = X1 - (i - 1) * ht_step
     Xc = X1 - (i - Morceau_STAiR_ds_dX_Pre) * ht_step
     Xb = X1 - i * ht_step

     If Xc > Xa Or Xc < Xb Then Stop

      'entre Xc et Xa Pds = P_startF * (1 - Exp(-(v - t_preSB) / to_SB))
      'entre Xb et Xc  Rds = P_startF * (1 - Exp(-(w - t_preSB) / to_SB))' 1 duree _Stair de plus pour le morceau_stair_ds_dX_pre

      Dds = IIf(Morceau_STAiR_ds_dX_Pre > 0 And drap = 1 And i = sN, P_startF, Rds)

      If F = 1 Then 'Fast NEW
         Qdp = Pdp * (1 - Exp(-Sdu_STAiR_V(i) / to_startF))
         Ddp = Dds * (1 - Exp(-Sdu_STAiR_V(i) / to_startF))
      Else 'Slow NEW
        Qdp = (P_startS + P_startF - Pdp) * (1 - Exp(-Sdu_STAiR_V(i) / to1_startS))
        Ddp = (P_startS + P_startF - Dds) * (1 - Exp(-Sdu_STAiR_V(i) / to1_startS))
      End If
If B = 1 And Icod = 1 And zp = sm And k = N_sm And i = sN Then Debug.Print "CALCUL ** step N° "; k, "F ="; F, "i = "; i, "xmi ="; Sdu_STAiR_V(i), "pT_avt = "; pT

      pT = Qdp * ((Xa - Xc) / dX_T) * (1 + 0.5 * (Xa + Xb) / Abs(X3))
      pT = pT + Ddp * ((Xc - Xb) / dX_T) * (1 + 0.5 * (Xa + Xb) / Abs(X3))

 If B = 1 And Icod = 1 And zp = sm And k = N_sm And i = sN Then Debug.Print "QDp ="; Qdp, "supp ="; Qdp * (ht_step / dX_T) * (1 + 0.5 * (Xa + Xb) / Abs(X3)), "pT = "; pT
      'If F = 2 And i = 2 And k = 8 Then Stop
```

```
      vd = "d" + Format(i)
      If Icod = 1 Or (Icod = 3 And F = 1 And IsNull(Escw(vd))) Then
        Escw(vd) = pT
      Else
        Escw(vd) = pT + Escw(vd)
      End If
      sY = sY + pT

  Next 'i

   If drap = 2 And Morceau_STAiR_ds_dX_Max > 0 Then 'pour l'instant je laisse tomner le cas ou morceau_stair_ds_dx_pre> morceau_stair ds dxmax et il faut rajouuet ce morceau mais c'est faisable
         'i = N_STAiR_ds_dX_Max+1
       Xa = X1 - (i - 1) * ht_step
       'If Morceau_STAiR_ds_dX_Pre > 0 And Morceau_STAiR_ds_dX_Max Then
       '   Xc = X3 - (i - Morceau_STAiR_ds_dX_Pre) * ht_step
       'Else
       '   Xc = X3
       'End If
       Xb = X3

       'If Xc > Xa Or Xc < Xb Then Stop

       'Dds = IIf(Morceau_STAiR_ds_dX_Pre > 0 And drap = 1 And i = N_F, P_startF, Rds)


      If F = 1 Then 'Fast NEW
        Qdp = Pdp * (1 - Exp(-Sdu_STAiR_V(i) / to_startF))
        'Ddp = Dds * (1 - Exp(-Sdu_STAiR_V(i) / to_startF))
      Else 'Slow NEW
         Qdp = (P_startS + P_startF - Pdp) * (1 - Exp(-Sdu_STAiR_V(i) / to1_startS))
        'Ddp = (P_startS + P_startF - Dds) * (1 - Exp(-Sdu_STAiR_V(i) / to1_startS))
      End If


          pT = Qdp * (Morceau_STAiR_ds_dX_Max * ht_step / dX_T) * (1 + 0.5 * (Xa + Xb) / Abs(X3))

         vd = "d" + Format(i)

         If Icod = 1 Or (Icod = 3 And F = 1 And IsNull(Escw(vd))) Then
            Escw(vd) = pT
         Else
            Escw(vd) = pT + Escw(vd)
         End If
         sY = sY + pT
   End If

End Sub
```

#### Sub AAPH_ESC_MON_calcul_pT1_elas_ISO_t_0(Icod As Integer)

```
'WS+ Rem : Icod= 1  1 valeur   Icd= 3 on somme
'pT1elas : Icod= 11 1 valeur   Icd= 13 on somme

'mise à 0 pour etre tranquille
For i = 1 To 15: Sdu_STAiR_V(i) = 0: Next
'faut inverser lstair 1 correspond à la durée k  etc
Sdu_STAiR_V(2) = du_STAiR_V(k - 1)

If k <= N_clik Then

  For i = (k - 2) To 1 Step -1
     TRUC = k - i + 1
     Sdu_STAiR_V(TRUC) = Sdu_STAiR_V(TRUC - 1) + du_STAiR_V(i)
  Next 'i

Else 'faut décaler en fonction de n_clik

  For i = (k - 2) To (k - N_clik) Step -1
     TRUC = k - i + 1
     Sdu_STAiR_V(TRUC) = Sdu_STAiR_V(TRUC - 1) + du_STAiR_V(i)
  Next 'i
End If

For i = N_F To sN

     Xa = X1 - (i - 1) * ht_step
     Xb = X1 - i * ht_step

     If F = 1 Then 'Fast en SB à t= 0
         Qdp = P_startF * (1 - Exp(-Sdu_STAiR_V(i) / to_startF))
     Else 'slow en SB à t=0
         Qdp = P_startS * (1 - Exp(-Sdu_STAiR_V(i) / to1_startS))
     End If
      pT = Qdp * (ht_step / dX_T) * (1 + 0.5 * (Xa + Xb) / Abs(X3))

      vd = "d" + Format(i)
      If Icod = 1 Or (Icod = 3 And F = 1) Then
        Escw(vd) = pT
      Else
        Escw(vd) = pT + Escw(vd)
      End If
      sY = sY + pT
 Next 'i

 If drap = 2 And Morceau_STAiR_ds_dX_Max > 0 Then
        If i <> (N_STAiR_ds_dX_Max + 1) Then Stop
        Xa = X1 - (i - 1) * ht_step
        Xb = X3

        If F = 1 Then 'Fast en SB à t= 0
            Qdp = P_startF * (1 - Exp(-Sdu_STAiR_V(i) / to_startF))
        Else 'slow en SB à t=0
          Qdp = P_startS * (1 - Exp(-Sdu_STAiR_V(i) / to1_startS))
        End If
        pT = Qdp * (Morceau_STAiR_ds_dX_Max * ht_step / dX_T) * (1 + 0.5 * (Xa + Xb) / Abs(X3))

        vd = "d" + Format(i)
       If Icod = 1 Or (Icod = 3 And F = 1) Then
          Escw(vd) = pT
        Else
          Escw(vd) = pT + Escw(vd)
        End If
        sY = sY + pT
  End If
  End Sub
```

#### Sub AAPH_ESC_MON_calcul_pT1_elas_NEW(Icod As Integer)

```
'WS+ Rem : Icod= 1  1 valeur   Icd= 3 on somme
'pT1elas : Icod= 11 1 valeur   Icd= 13 on somme
```

```
If N_F > sN Then Stop '?

'faut inverser lstair 1 correspond à la durée k  etc
Sdu_STAiR_V(2) = du_STAiR_V(k - 1)

If k <= N_clik Then

  For i = (k - 2) To 1 Step -1
     TRUC = k - i + 1
     Sdu_STAiR_V(TRUC) = Sdu_STAiR_V(TRUC - 1) + du_STAiR_V(i)
  Next 'i

Else 'faut décaler en fonction de n_clik

  For i = (k - 2) To (k - N_clik) Step -1
     TRUC = k - i + 1
     Sdu_STAiR_V(TRUC) = Sdu_STAiR_V(TRUC - 1) + du_STAiR_V(i)
  Next 'i
End If

'pre
Rds = 0
If to2_SB = 0 Then 'simple exponentielle
  Pdp = P_startF * (1 - Exp(-(Sdu_STAiR_V(N_STAiR_ds_dX_Pre + 1) - t_preSB) / to1_SB))
  If Morceau_STAiR_ds_dX_Pre > 0 Then
   Rds = P_startF * (1 - Exp(-(Sdu_STAiR_V(N_STAiR_ds_dX_Pre + 2) - t_preSB) / to1_SB)) 'Morceau ds dX_pre
  End If

Else '* ** double exponentizelles
    sX = to1_SB * Exp(-(Sdu_STAiR_V(N_STAiR_ds_dX_Pre + 1) - t_preSB) / to1_SB) / (to1_SB - to2_SB)
    sXX = to2_SB * Exp(-(Sdu_STAiR_V(N_STAiR_ds_dX_Pre + 1) - t_preSB) / to2_SB) / (to2_SB - to1_SB)
    Pdp = P_startF * (1 - sX - sXX)
  If Morceau_STAiR_ds_dX_Pre > 0 Then
     sX = to1_SB * Exp(-(Sdu_STAiR_V(N_STAiR_ds_dX_Pre + 2) - t_preSB) / to1_SB) / (to1_SB - to2_SB) 'Morceau ds dX_pre
     sXX = to2_SB * Exp(-(Sdu_STAiR_V(N_STAiR_ds_dX_Pre + 2) - t_preSB) / to2_SB) / (to2_SB - to1_SB) 'Morceau ds dX_pre
     Rds = P_startF * (1 - sX - sXX)
  End If
End If

For i = N_F To sN
     ' STAiR = [Xb Xc]_RDS + (Xc ; Xa]_PdP
     Xa = X1 - (i - 1) * ht_step
     Xc = X1 - (i - Morceau_STAiR_ds_dX_Pre) * ht_step
     Xb = X1 - i * ht_step

     If Xc > Xa Or Xc < Xb Then Stop

      'entre Xc et Xa Pds = P_startF * (1 - Exp(-(v - t_preSB) / to_SB))
      'entre Xb et Xc  Rds = P_startF * (1 - Exp(-(w - t_preSB) / to_SB))' 1 duree _Stair de plus pour le morceau_stair_ds_dX_pre

      Dds = IIf(Morceau_STAiR_ds_dX_Pre > 0 And drap = 1 And i = sN, P_startF, Rds)


      If F = 1 Then 'Fast NEW
         Qdp = Pdp * (1 - Exp(-Sdu_STAiR_V(i) / to_startF))
         Ddp = Dds * (1 - Exp(-Sdu_STAiR_V(i) / to_startF))
      Else 'Slow NEW
         Qdp = (P_startS + P_startF - Pdp) * (1 - Exp(-Sdu_STAiR_V(i) / to1_startS))
         Ddp = (P_startS + P_startF - Dds) * (1 - Exp(-Sdu_STAiR_V(i) / to1_startS))
      End If
      pT = Qdp * ((Xa - Xc) / dX_T) * (1 + 0.5 * (Xa + Xb) / Abs(X3))
      pT = pT + Ddp * ((Xc - Xb) / dX_T) * (1 + 0.5 * (Xa + Xb) / Abs(X3))

      vd = "d" + Format(i)
      If Icod = 1 Or (Icod = 3 And F = 1 And IsNull(Escw(vd))) Then
         Escw(vd) = pT
      Else
         Escw(vd) = pT + Escw(vd)
      End If
      sY = sY + pT
```

```
  Next 'i

   If drap = 2 And Morceau_STAiR_ds_dX_Max > 0 Then 'pour l'instant je laisse tomner le cas ou morceau_stair_ds_dx_pre> morceau_stair ds dxmax et il faut rajouuet ce morceau mais c'est faisable
          'i = N_STAiR_ds_dX_Max+1
        Xa = X1 - (i - 1) * ht_step
        'If Morceau_STAiR_ds_dX_Pre > 0 And Morceau_STAiR_ds_dX_Max Then
        '   Xc = X3 - (i - Morceau_STAiR_ds_dX_Pre) * ht_step
        'Else
        '   Xc = X3
        'End If
        Xb = X3

        'If Xc > Xa Or Xc < Xb Then Stop

        'Dds = IIf(Morceau_STAiR_ds_dX_Pre > 0 And drap = 1 And i = N_F, P_startF, Rds)


       If F = 1 Then 'Fast NEW
          Qdp = Pdp * (1 - Exp(-Sdu_STAiR_V(i) / to_startF))
          'Ddp = Dds * (1 - Exp(-Sdu_STAiR_V(i) / to_startF))
       Else 'Slow NEW
          Qdp = (P_startS + P_startF - Pdp) * (1 - Exp(-Sdu_STAiR_V(i) / to1_startS))
          'Ddp = (P_startS + P_startF - Dds) * (1 - Exp(-Sdu_STAiR_V(i) / to1_startS))
       End If


          pT = Qdp * (Morceau_STAiR_ds_dX_Max * ht_step / dX_T) * (1 + 0.5 * (Xa + Xb) / Abs(X3))

          vd = "d" + Format(i)

          If Icod = 1 Or (Icod = 3 And F = 1 And IsNull(Escw(vd))) Then
             Escw(vd) = pT
          Else
             Escw(vd) = pT + Escw(vd)
          End If
          sY = sY + pT
   End If

End Sub
```
