## Supplementary Chapter. Theoretical study of a staircase shortening for "Mechanical model of muscle contraction. 6. Calculations of the tension exerted by a skeletal fiber during a shortening staircase"

### S6.L Supplement Chapter of Paper 6.

### Theoretical study of a staircase shortening.

#### L.1 Staircase step n° i included in the linear domain

The angle θ defines the position of the lever (S1b) belonging to a myosin II head in working stroke (WS). To characterize the linear displacement of a half-sarcomere (hs) along the longitudinal axis of the myofibril, the X abscissa of the embedding point of the motor domain (S1a) on the actin filament is used relative to the point representing the pivot connection between the rod (S2) and the myosin filament. The two parameters, θ and X, are linked by an affine function formulated in (I21a) in Supplement S4.I of accompanying Paper 4, reproduced below:

$$X = L_{S1b} \cdot R_{WS} \cdot (\theta - \theta_0) \tag{L1}$$

where $L_{S1b}$ is the length of the lever S1b; $R_{WS}$ is a geometric constant equal to about 0.95 determined in equalities (13) and (14) from Paper 2; $\theta_0$ is the middle of the angular range $\delta\theta_T$ calculated with formula (4) in Paper 4.

Equation (L1) determines a linear domain bounded by the angles $\theta_{down}$ and $\theta_{up}$ (Fig L1a) and by the corresponding abscissa $X_{down}$ and $X_{up}$ (Fig I1 of Supplement S4.I). With respect to step n° i of a staircase shortening, the iterative displacement "$\Delta X_i = -\delta X_{stair}$" leads to iterative rotation "$\Delta\theta_i = -\delta\theta_{stair}$" in the linear domain. The angular range $\delta\theta_{stair}$ is associated to the linear range $\delta X_{stair}$ according to (L1):

$$\delta\theta_{stair} = \frac{\delta X_{stair}}{L_{S1b} \cdot R_{WS}} \tag{L2}$$

#### L.2 Instantaneous proportion of heads that can quickly initiate a WS (recall)

The {startF} event refers to the Fast initiation of a WS and requires the Strong Binding (SB) state where the myosin head is strongly bound. The probability ($P_{SB}$) of occurrence of the event {SB} is given in (B11) in paragraph B.4 of Supplement S1.B of Paper 1 and reproduced below:

$$P_{SB}(t) = p_{startF} \cdot \left(1 - e^{-\frac{t - \tau_{preSB}}{\tau_{SB}}}\right) \tag{L3}$$

where $p_{startF}$ is the maximum proportion of heads capable of rapidly initiating a WS after a shortening following the isometric tetanus plateau; $p_{startF}$ is represented by a dark blue rectangle with grid of black dots arranged diagonally in Fig L1a.

**Note:** the isolated event {SB} only concerns the event {startF} because the SB state and its related transitions are integrated in the events {startS} and {startVS}, global events related to the Slow initiation and the Very Slow initiation of a WS; see Supplement S1.B relative to Paper 1.

#### L.3 Step-by-step description of a staircase shortening

The sequence of a staircase shortening from the tetanus plateau to the repetitive regime is described below and is illustrated with the graphs in Figs L1a to L1g.

**Isometric tetanus plateau (Fig L1a)**

The tension of the isometric tetanus plateau (T0) is the subject of paragraph I.3 of Supplement S4.I relative to Paper 4. Resulting from the global action of the $\Lambda_0$ WS heads in each hs, T0 is the sum of the linking forces induced by the $\Lambda_0$ motor-moments applied to the $\Lambda_0$ levers S1b whose angular positions $\theta$ are uniformly distributed between the angles $\theta_T$ and $\theta_{up}$; see green rectangle in Fig L1a, the grid with black dots aligned horizontally indicating the presence of heads in WS state during the isometric tetanus plateau preceding the first step.

To the right of the $\theta_{up}$ terminal appears a dark blue rectangle whose width is equal to $\delta\theta_{pre}$. This rectangle quoted in the previous paragraph represents the uniform distribution of SB heads during the isometric tetanus plateau, corresponding to $p_{startF}$, the maximum proportion of heads likely to be in SB. This display is educational because in reality we do not know if the lever belonging to a SB head is located in the fixed plane containing the longitudinal axis of the hs, a plane defined with hypothesis 4 characterizing the WS state (see accompanying Paper 2). The model simulates a uniform distribution of the $\theta$ angle because after the hs shortening equal to $-\delta X_{stair}$, a myosin head is able to quickly initiate a WS if its lever has an angular position $\theta$ corresponding to the rotation $-\delta\theta_{stair}$ in the linear domain. Thus all the distributions to the right of $\theta_{up}$ represented by light or dark blue rectangles in the different schemes of the Fig L1 are, to date, fictitious and are only the transposed images of the uniform distributions located, in the linear domain, to the left of $\theta_{up}$ in a hs on the right (to the right of $\theta_{up}$ in a hs on the left).

**Step 1 (Fig L1b): the hs is shortened by "$\Delta X_1 = - \delta X_{stair}$"**

At $t=\tau_{p1}$, i.e. at the end of phase 1 of the first step, the levers of myosin heads previously in WS state have rotated by "$\Delta\theta_1 = - \delta\theta_{stair}$" in accordance with equality (L2) and their $\theta$ angles are uniformly distributed between ($\theta_T - \delta\theta_{stair}$) and ($\theta_{up} - \delta\theta_{stair}$); the density of the $\theta$ angle is represented by a green rectangle with horizontal dot grid. No WS initiation is possible in this area.

In the angular range $\delta\theta_{stair}$ released between the two terminals ($\theta_{up} - \delta\theta_{stair}$) and $\theta_{up}$, myosin heads strongly linked to t=0 quickly initiate a WS with the event {startF} (yellow rectangle) and non-SB heads at t=0 slowly initiate a WS with the event {startS} if $\tau_{stair} \leq t < \tau_{preS}$ (pink rectangle). The grid of black dots arranged diagonally into the pink rectangle indicates the presence of heads in SB state during the isometric tetanus plateau, i.e. before the first staircase step.

To the left of $\theta_{up}$ in the area freed by the rotation $\Delta\theta_1$, myosin heads initiate according to (L3) a SB (light blue rectangle without dot grid).

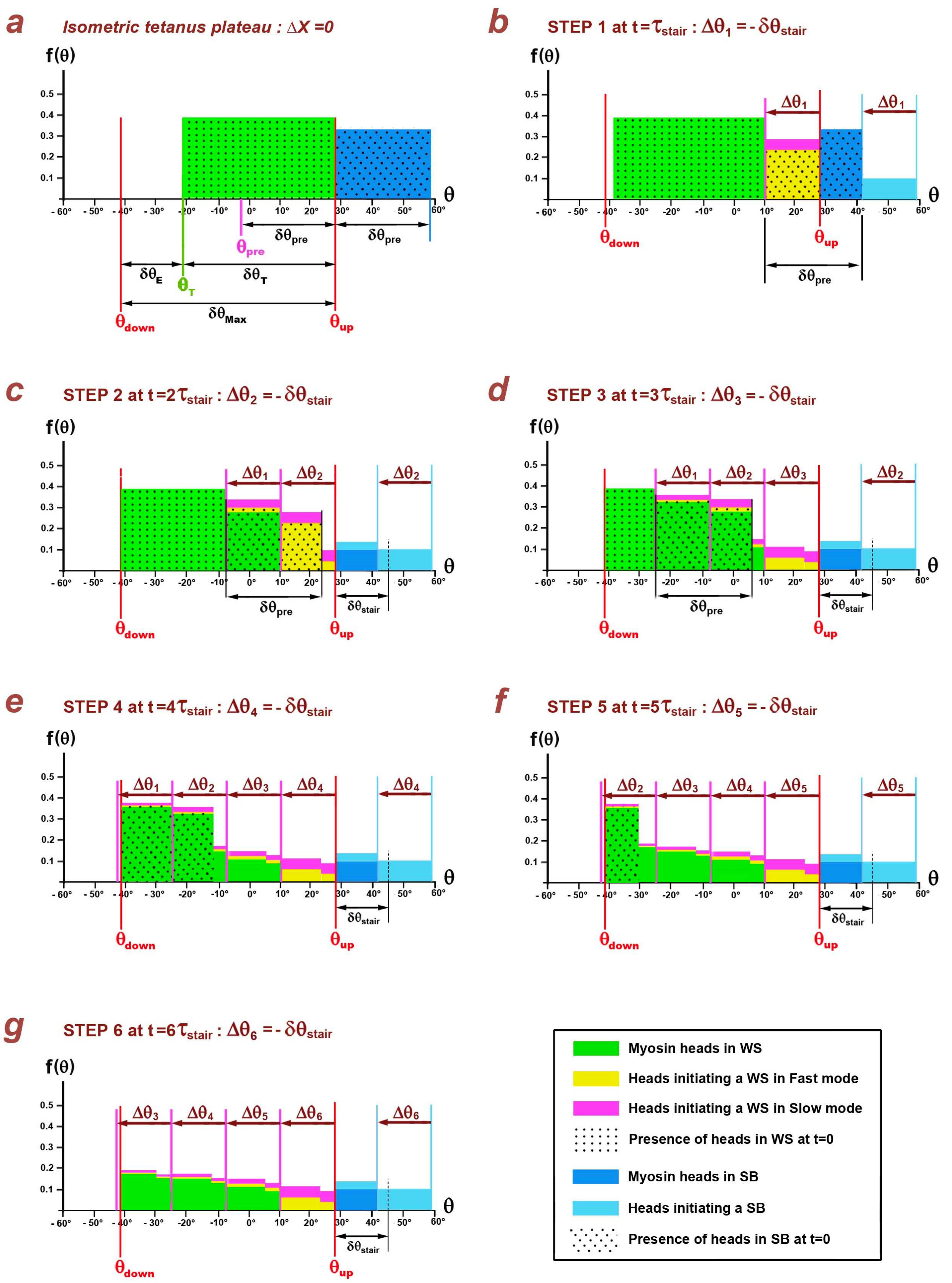


**Fig L1. Uniform distributions of the θ angle in a half-sarcomere on the right during a shortening staircase.**

(a) Isometric tetanus plateau. (b) to (g) From the first staircase step to the repetitive regime.

A $t=\tau_{stair}$, i.e. at the time end of step 1, the tension is calculated by assessing each of the contributions, zone by zone in the linear domain, taking into account the slowly detaching heads whose levers have an orientation $\theta$ between $\theta_{down}$ and $\theta_T$. Heads that detach quickly with the {FastDE} event are excluded because their number is negligible. In addition, we omit the very slow initiation of the WS because the duration of a staircase step is in practice much lower than the occurrence delay of the event {startVS}, i.e. $\tau_{stair} << \tau_{preVS}$.

**Step 2 (Fig L1c): the hs is shortened by "$\Delta X_2 = - \delta X_{stair}$"**

At $t=(\tau_{stair}+\tau_{p1})$, i.e. at the end of phase 1 relating to the second step, all levers of heads previously in WS to $t=\tau_{stair}$ have rotated by "$\Delta\theta_2 = - \delta\theta_{stair}$" in accordance with the equality (L2).

The angular range released between $(\theta_{up} - \delta\theta_{stair})$ and $\theta_{up}$ is divided into two parts:

1/ between $(\theta_{up} - \delta\theta_{stair})$ and $(\theta_{up} – [2\cdot\delta\theta_{stair} - \delta\theta_{pre}])$: SB heads at t=0 initiate quickly (yellow rectangle with diagonal dot grid) and non-SB heads at t=0 slowly initiate a WS (pink rectangle).

2/ between $(\theta_{up} – [2\cdot\delta\theta_{stair} - \delta\theta_{pre}])$ and $\theta_{up}$: heads that have had time to bind strongly according to {SB} event during $\tau_{stair}$, i.e. the duration of step 1, quickly initiate a WS (yellow rectangle without dot grid) with a lower proportion compared to that of the first part since the proportion of heads previously in SB is lower than $p_{startF}$ according to (L3). Non-SB heads slowly initiate a WS (pink rectangle).

In the range between the two terminals $(\theta_{up} - 2\cdot\delta\theta_{stair})$ and $(\theta_{up} - \delta\theta_{stair})$, we find the portion of heads that have initiated a WS (green rectangle with diagonal dot grid). Heads continue to initiate a WS quickly (yellow rectangle with diagonal dot grid).and slowly (pink rectangle).

Within the range between the two terminals $(\theta_{up} - 2\cdot\delta\theta_{stair})$ and $(\theta_{up} - \delta\theta_{stair})$, there remains the portion of WS heads at t=0 (green rectangle with horizontal dot grid). No WS initiation is possible in this area.

To the left of $\theta_{up}$ in the area freed by the rotation $\Delta\theta_2$, heads initiate according to (L3) a strong binding (light blue rectangle without dot grid) and in the second extension area $(\theta_{pre} - \delta\theta_{stair})$, we find the SB heads at step 1 (dark blue rectangle) and new heads initiate according to (L3) a strong binding (light blue rectangle). It should be noted that no more dot grid are present and that the overall pattern on $\delta\theta_{pre}$ is identical in the following graphs: the repetitive regime with respect to the {SB} event is in place.

At $t=2\cdot\tau_{stair}$, i.e. at the time end of step 2, the tension is calculated by taking stock of each of the contributions, zone by zone in the linear domain.

**Step 3 (Fig L1d): the hs is shortened by "$\Delta X_3$ = - $\delta X_{stair}$ "**

At t=($2 \cdot \tau_{stair}+\tau_{p1}$), i.e. at the end of phase 1 of the third step, all WS head levers at t = $2 \cdot \tau_{stair}$ have rotated by "$\Delta\theta_3$ = - $\delta\theta_{stair}$".

In the space freed between the two terminals ($\theta_{up}$ - $\delta\theta_{stair}$) and $\theta_{up}$, previously SB heads initiate a WS rapidly (yellow rectangles) and non-SB heads slowly initiate a WS (pink rectangles): the reproducibility regime is operational for this range which is unchanged in all the following figures.

In the range between the two terminals ($\theta_{up}$ - $2 \cdot \delta\theta_{stair}$) and ($\theta_{up}$ - $\delta\theta_{stair}$) we recognize the portion of heads that initiated a WS to the previous step in the neighbouring area (green rectangles with and without diagonal dot grid). Heads continue to initiate a WS rapidly (yellow rectangles with and without diagonal dot grid) and slowly (pink rectangles).

Within the range between the two terminals ($\theta_{up}$ - $3 \cdot \delta\theta_{stair}$) and ($\theta_{up}$ - $2 \cdot \delta\theta_{stair}$), we can distinguish the portion of heads that initiated a WS to the previous step in the neighbouring area (green rectangle with diagonal dot grid). Heads continue to initiate a WS rapidly (yellow rectangle with diagonal dot grid) and slowly (pink rectangle).

Within the range between the two terminals $\theta_{down}$ and ($\theta_{up}$ - $3 \cdot \delta\theta_{stair}$), there remains the portion of WS heads at t=0 (green rectangle with horizontal dot grid). No WS initiation is possible in this area.

At t=$3 \cdot \tau_{stair}$, i.e. at the time end of step 3, the tension is calculated by taking stock of each of the contributions, area by area in the linear domain.

**Stairs 4, 5 and 6 (Figs L1e, L1f and L1g)**

The different processes described for the previous steps are implemented at each shortening "$\Delta X_i$ = - $\delta X_{stair}$", until step 6 where all the dot grids have disappeared (Fig L1g). The identical reproducibility operates for the following steps: the regime of the heads in WS or slowly detaching is repetitive.

At the time end of step 6, the tension is calculated by taking stock of each of the contributions, area by area. This tension is the constant repetitive tension called as T*.

#### L. 4 Calculation of the tension generated by the WS heads in repetitive regime

##### *L.4.1 Number of steps required to reach the repetitive scheme (n*)*

We characterize the event E:

$$E \equiv \left\{ \left( \frac{\delta\theta_{pre} + \delta\theta_{Max}}{\delta\theta_{stair}} \right) > \mathrm{int}\left( \frac{\delta\theta_{pre} + \delta\theta_{Max}}{\delta\theta_{stair}} \right) \right\}$$

where int is the integer part symbol.

The repetitive regime occurs at step n*, when all the heads "prepared" in state SB and all the heads initially in state WS at t=0 have disappeared, that is:

$$n^* = \left[ \mathrm{int}\left( \frac{\delta\theta_{pre} + \delta\theta_{Max}}{\delta\theta_{stair}} \right) + \mathbf{1}_E \right] \qquad \text{(L4)}$$

where $\mathbf{1}$ is the indicator function defined in (A2a) in Supplement S1.A of Paper 1.

##### *L.4.2 Segmentation of linear ranges according to $\delta X_{stair}$*

The two linear extensions $\delta X_E$ and $\delta X_{Max}$ are introduced in (I23) and (I24) in Supplement S4.I of accompanying Paper 4; see Fig I1. The linear range $\delta X_{pre}$ which corresponds according to (L1) to the angular range $\delta\theta_{pre}$ (Fig L1) is presented in the Methods section of Paper 1.

A value of $\delta X_{stair}$ is chosen small enough that each of these three ranges is a multiple of $\delta X_{stair}$ such that:

$$\delta X_{Max} = n_M \cdot \delta X_{stair} \qquad \text{(L5a)}$$

$$\delta X_E = n_E \cdot \delta X_{stair} \qquad \text{(L5b)}$$

$$\delta X_{pre} = n_{pre} \cdot \delta X_{stair} \qquad \text{(L5c)}$$

where $n_M$, $n_E$ and $n_{pre}$ are three integers greater than 1.

These three relationships are translated into angular ranges with (L1):

$$\delta\theta_{Max} = n_M \cdot \delta\theta_{stair} \qquad \text{(L6a)}$$

$$\delta\theta_E = n_E \cdot \delta\theta_{stair} \qquad \text{(L6b)}$$

$$\delta\theta_{pre} = n_{pre} \cdot \delta\theta_{stair} \qquad \text{(L6c)}$$

Note: in the example in Fig L1, and in the study cases of physiologic literature, the segmentation of $\delta X_E$, $\delta X_{Max}$ and $\delta X_{pre}$ by $\delta X_{stair}$ are not exact; our calculation algorithms take into account the remains of the divisions.

##### *L.4.3 Segmentation of durations according to $\tau_{stair}$*

The {startVS} event concerning the very slow initiation of a WS has been neglected in the description given in paragraph L.3; we reintroduce this event for the developments of paragraphs L.5 and L.7 devoted to the transition to continuous.

The delays $\tau_{preS}$ and $\tau_{preVS}$ for the implementation of {startS} and {startVS} events are provided in expressions (B5b) and (B5c) of Supplement S1.B of Paper 1. The occurrence delay $\tau_{preSDE}$ related to the slow detachment of a WS head, i.e. the event {SlowDE} appears in (B13). We choose a value of $\tau_{stair}$ short enough so that $\tau_{preS}$, $\tau_{preVS}$ and $\tau_{preSDE}$ are each a multiple of $\tau_{stair}$:

$$\tau_{preS} = n_{preS} \cdot \tau_{stair} \tag{L7a}$$

$$\tau_{preVS} = n_{preVS} \cdot \tau_{stair} \tag{L7b}$$

$$\tau_{preSDE} = n_{preSDE} \cdot \tau_{stair} \tag{L7c}$$

where $n_{preS}$ , $n_{preVS}$ and $n_{preSDE}$ are three integers greater than 1.

##### *L.4.4 Maximum proportions of heads that can initiate a WS in repetitive mode*

**Fast initiation with {startF}**: in repetitive regime at the time end of the staircase step, the maximum proportion of heads that can initiate a WS in fast mode ($P^*_{SB}$) is calculated with (L3) and (L5c) according to:

$$P^*_{SB} = P_{SB}(n_{pre} \cdot \tau_{stair}) = p_{startF} \cdot \left(1 - e^{-\frac{(n_{pre} \cdot \tau_{stair} - \tau_{preSB})}{\tau_{SB}}}\right) \tag{L8a}$$

The proportion of heads likely to transit in SB state and who did not have time to achieve the event {SB} at the time end of the step is:

$$P^*_{\overline{SB}} = p_{startF} - P^*_{SB}$$

**Slow initiation with {startS}**: the maximum proportion of heads slowly initiating a WS in repetitive mode ($P^*_{Slow}$) is equal to the maximum proportion of heads able to initiate a WS in slow mode after a shortening following isometric tetanus plateau, a proportion named "$p_{startS}$" and defined in (B4b) in Supplement S1.B of Paper 1, to which must be added the proportion of heads likely to transit in SB state without performing it:

$$P^*_{Slow} = \left(p_{startS} + P^*_{\overline{SB}}\right) \cdot \mathbb{1}_{[\tau_{preS};+\infty[}(t)$$

So:

$$P^*_{Slow} = \left(p_{startF} + p_{startS} - P^*_{SB}\right) \cdot \mathbb{1}_{[\tau_{preS};+\infty[}(t) \tag{L8b}$$

**Very slow initiation with {startVS}** : the maximum proportion of heads initiating very slowly a WS in repetitive regime ($P^*_{VSlow}$) is equal to the maximum proportion of heads able to initiate a WS in very slow mode after a length step following the isometric tetanus plateau, proportion named "$p_{startVS}$" defined in (B4c) and equal to (1- $p_{startF}$ - $p_{startS}$) with (B4d), so:

$$P^*_{VSlow} = \left(1 - p_{startF} - p_{startS}\right) \cdot \mathbb{1}_{[\tau_{preVS};+\infty[}(t) \tag{L8c}$$

##### *L.4.5 Instantaneous proportions and numbers of heads initiating a WS in repetitive regime*

In paragraph B.3 of Supplement S1.B of Paper 1, the global event {WSstart} leading to the WS state is declined into three disjointed modes, fast with {startF}, slow with {startS} or very slow with {startVS}, events whose time equations are presented in (B5a), (B5b) and (B5c), respectively.

The two events {SB} and {startF} are independent since a mechanical movement is necessary for {startF} to occur. The instantaneous proportion of heads initiating a WS rapidly in repetitive mode ($P^*_F$) is equal to the product of their 2 probabilities:

$$P^*_F(t) = P^*_{SB} \cdot \left(1 - e^{-\frac{t}{\tau_{startF}}}\right) \tag{L9a}$$

where $P^*_{SB}$ is given in (L8a).

The instantaneous proportion of heads slowly initiating a WS in repetitive mode ($P^*_S$) is formulated with (L7a):

$$P^*_S(t) = P^*_{Slow} \cdot \left(1 - e^{-\frac{t - n_{preS} \cdot \tau_{stair}}{\tau_{startS}}}\right) \tag{L9b}$$

where $P^*_{Slow}$ is given in (L8b).

The instantaneous proportion of heads initiating very slowly a WS in repetitive mode ($P^*_{VS}$) is written with (L7b):

$$P^*_{VS}(t) = P^*_{VSlow} \cdot \left(1 - e^{-\frac{t - n_{preVS} \cdot \tau_{stair}}{\tau_{startVS}}}\right) \tag{L9c}$$

where $P^*_{VSlow}$ is given in (L8c).

In total:

$$P^*_{WS}(t) = P^*_F(t) + P^*_S(t) + P^*_{VS}(t) \tag{L10}$$

We consider the staircase step n° i with i ≥ n*. Since the realization of a WS occurs only on the range $\delta\theta_{Max}$ between $\theta_{down}$ and $\theta_{up}$, the number of steps involved in the calculation of the tension is equal to $n_M$ according to (L5a) and (L6a).

The total number of WS heads per hs ($\Lambda^*_{WS}$) at the time end of step n° i in repetitive mode is calculated by summing the expression (L10) for each of the $n_M$ indexed steps from i to [i - ($n_M$ -1)], and by using the equality (I11) of the Supplement S4.I of Paper 4. So, we got by affine transformation with (L1):

$$\overset{*}{\Lambda}_{WS}(i\cdot\tau_{sair}) = \left(\Lambda_0\cdot\frac{\delta X_{stair}}{\delta X_T}\right)\cdot\left(\sum_{k=1}^{n_M}\left[\overset{*}{P}_{WS}(k\cdot\tau_{stair})\right]\right) \tag{L11}$$

where step n° i corresponds to k=1, step n° (i-1) to k=2, step n° (i-2) to k=3, ..., until step n°[i-($n_M$-1)] which corresponds to k=$n_M$; $\Lambda_0$ is the number of WS heads per hs during the isometric tetanus plateau.

In repetitive regime, the number of heads in WS depends on $\delta X_{stair}$ and $\tau_{stair}$ but is independent of the index i.

The expression (L11) is rewritten with (L10), and more specifically with the equalities (L7a) and (L7b) relating to the two events {startS} and {startVS}:

$$\overset{*}{\Lambda}_{WS}(i\cdot\tau_{sair}) = \overset{*}{\Lambda}_{F}(i\cdot\tau_{sair}) + \overset{*}{\Lambda}_{S}(i\cdot\tau_{sair}) + \overset{*}{\Lambda}_{VS}(i\cdot\tau_{sair}) \tag{L12}$$

with

$$\overset{*}{\Lambda}_{F}(i\cdot\tau_{sair}) = \left(\Lambda_0\cdot\frac{\delta X_{stair}}{\delta X_T}\right)\cdot\sum_{k=1}^{n_M}\left[\overset{*}{P}_{F}(k\cdot\tau_{stair})\right] \tag{L13a}$$

$$\overset{*}{\Lambda}_{S}(i\cdot\tau_{sair}) = \left(\Lambda_0\cdot\frac{\delta X_{stair}}{\delta X_T}\right)\cdot\sum_{k=(n_{preS}+1)}^{n_M}\left[\overset{*}{P}_{S}(k\cdot\tau_{stair})\right] \tag{L13b}$$

$$\overset{*}{\Lambda}_{VS}(i\cdot\tau_{sair}) = \left(\Lambda_0\cdot\frac{\delta X_{stair}}{\delta X_T}\right)\cdot\sum_{k=(n_{preVS}+1)}^{n_M}\left[\overset{*}{P}_{VS}(k\cdot\tau_{stair})\right] \tag{L13c}$$

##### *L.4.6 Tension generated by WS heads in repetitive regime*

The calculation for a single step is carried out at the time end of staircase step n° [i-(k-1)] corresponding to the k value.

**Conditions with (L10):**

$$\left.\begin{array}{r} p = \overset{*}{P}_{WS}(k\cdot\tau_{stair}) \\ X_1 = X_{up} - (k-1)\cdot\delta X_{stair} \\ X_2 = X_{up} - k\cdot\delta X_{stair} \end{array}\right\} \Rightarrow \begin{array}{c} \delta X_L = X_1 - X_2 = \delta X_{stair} \\ \\ X_1 + X_2 = 2\cdot X_{up} - (2k-1)\cdot\delta X_{stair} \end{array}$$

**Application with equation (I31b) displayed in Supplement S4.I of Paper 4:**

$$pT_{stair} = \overset{*}{P}_{WS}(k\cdot\tau_{stair})\cdot\left(\frac{\delta X_{stair}}{\delta X_T}\right)\cdot\frac{(\delta X_{Max} - (k-1/2)\cdot\delta X_{stair})}{|X_{down}|}$$

By summing the tensions relative to each of the $n_M$ steps, the total relative tension generated by the WS heads $\overset{*}{\Lambda}_{WS}(i\cdot\tau_{sair})$ at the time end of step i (with i ≥ n*) in repetitive regime is:

$$p\overset{*}{T}_{WS}(i\cdot\tau_{stair}) = \left(\frac{\delta X_{stair}}{\delta X_T}\right)\cdot\sum_{k=1}^{n_M}\left[\overset{*}{P}_{WS}(k\cdot\tau_{stair})\cdot\frac{(\delta X_{Max} - (k-1/2)\cdot\delta X_{stair})}{|X_{down}|}\right] \quad \text{(L14)}$$

In repetitive mode, the tension depends on $\delta X_{stair}$ and $\tau_{stair}$ but is independent of the index i as provided for observations.

Equality (L14) is rewritten with (L10):

$$p\overset{*}{T}_{WS}(i\cdot\tau_{sair}) = p\overset{*}{T}_F(i\cdot\tau_{sair}) + p\overset{*}{T}_S(i\cdot\tau_{sair}) + p\overset{*}{T}_{VS}(i\cdot\tau_{sair}) \quad \text{(L15)}$$

with

$$p\overset{*}{T}_F(i\cdot\tau_{sair}) = \left(\frac{\delta X_{stair}}{\delta X_T}\right)\cdot\sum_{k=1}^{n_M}\left[\overset{*}{P}_F(k\cdot\tau_{stair})\cdot\frac{(\delta X_{Max} - (k-1/2)\cdot\delta X_{stair})}{|X_{down}|}\right] \quad \text{(L16a)}$$

$$p\overset{*}{T}_S(i\cdot\tau_{sair}) = \left(\frac{\delta X_{stair}}{\delta X_T}\right)\cdot\sum_{k=(n_{preS}+1)}^{n_M}\left[\overset{*}{P}_S(k\cdot\tau_{stair})\cdot\frac{(\delta X_{Max} - (k-1/2)\cdot\delta X_{stair})}{|X_{down}|}\right] \quad \text{(L16b)}$$

$$p\overset{*}{T}_{VS}(i\cdot\tau_{sair}) = \left(\frac{\delta X_{stair}}{\delta X_T}\right)\cdot\sum_{k=(n_{preVS}+1)}^{n_M}\left[\overset{*}{P}_{VS}(k\cdot\tau_{stair})\cdot\frac{(\delta X_{Max} - (k-1/2)\cdot\delta X_{stair})}{|X_{down}|}\right] \quad \text{(L16c)}$$

#### L. 5 Calculation of the tension generated by the WS heads when a hs is shortening to steady velocity

##### *L.5.1 Constant speed (u) of hs shortening*

The two parameters characterizing a step, length ($\delta X_{stair}$) and duration ($\tau_{stair}$), are gradually decreased until they tend towards infinitesimal values:

$$\Delta X = -\delta X_{stair} \quad \rightarrow \quad dX \tag{L17a}$$

$$\tau_{stair} \quad \rightarrow \quad dt \tag{L17b}$$

Thus we approach a shortening carried out at a constant speed (u) for each hs such that:

$$u = -\frac{\delta X_{stair}}{\tau_{stair}} = \frac{dX}{dt} \tag{L18}$$

**Note**: u is expressed in algebraic value and therefore negative for a shortening (dX<0).

By integrating (L18) into the linear domain, we obtain with $X(t=0)=X_{up}$ :

$$\int_{X_{up}}^{X} dX = u \cdot \int_{0}^{t} dt$$

So:

$$(X - X_{up}) = u \cdot t \tag{L19}$$

The abscissa $X_S$ and $X_{VS}$ relating to the occurrence delays of the two events {startS} and {startVS} are defined using (L7a), (L7b) and (L19) as:

$$X_S = X_{up} + u \cdot \tau_{preS} \tag{L20a}$$

$$X_{VS} = X_{up} + u \cdot \tau_{preVS} \tag{L20b}$$

##### *L.5.2 Maximum proportions of heads that can initiate a WS when the hs is shortened at constant speed u*

With (L5c) and (L19), the following equality is checked in the linear domain:

$$n_{pre} \cdot \tau_{stair} = -\frac{\delta X_{pre}}{u} \tag{L21}$$

When shortening at steady velocity, the maximum proportion of heads that can initiate a WS quickly in repetitive mode ($P^*_{SB}$) tends towards a proportion ($P_F$) function of u that is calculated from (L8a) with (L21) according to:

$$P^*_{SB} \rightarrow P_F(u) = p_{startF} \cdot \left(1 - e^{\frac{\delta X_{pre} + u \cdot \tau_{preSB}}{u \cdot \tau_{SB}}}\right) \tag{L22a}$$

The maximum proportion of heads that can initiate a WS slowly in repetitive mode ($P^*_{Slow}$) tends towards a proportion ($P_S$) function of u that is calculated from (L8b), (L20a) and (L22a):

$$P^*_{Slow} \rightarrow P_S(u) = \left[p_{startS} + p_{startF} - P_F(u)\right] \cdot \mathbf{1}_{\{|u| \cdot \tau_{preS}) \leq \delta X_{Max}\}}(u) \qquad \text{(L22b)}$$

The maximum proportion of heads that can initiate a WS very slowly in repetitive mode ($P^*_{VSlow}$) tends towards a proportion ($P_{VS}$) function of u that is calculated from (L8c) and (L20b) according to:

$$P^*_{VSlow} \rightarrow P_{VS}(u) = \left[1 - p_{startF} - p_{startS}\right] \cdot \mathbf{1}_{\{|u| \cdot \tau_{preVS}) \leq \delta X_{Max}\}}(u) \qquad \text{(L22c)}$$

##### *L.5.3 Numbers and tensions generated by WS heads initiated according to {startF} when hs is shortened at constant speed u*

**Number of WS heads having initiated according to {startF}**

With (L9a), the expression (L13a) is reformulated:

$$\Lambda^*_F(i \cdot \tau_{sair}) = \left(\frac{\Lambda_0 \cdot P^*_{SB}}{\delta X_T}\right) \cdot \sum_{k=1}^{n_M} \left[\left(1 - e^{-\frac{k \cdot \tau_{stair}}{\tau_{startF}}}\right) \cdot \delta X_{stair}\right]$$

We apply (L17a) and (L17b) to approach a constant velocity of shortening: the factor ($k \cdot \tau_{stair}$) tends towards t and the number of heads ($\Lambda^*_F$) quickly initiating a WS on $\delta X_{Max}$, i.e. between $X_{up}$ and $X_{down}$, tends towards a number ($\Lambda_F$), function of u, which is calculated with (L22a) according to:

$$\Lambda^*_F \rightarrow \Lambda_F(u) = \left(\frac{\Lambda_0 \cdot P_F(u)}{\delta X_T}\right) \cdot \int_{X_{down}}^{X_{up}} \left(1 - e^{-\frac{t}{\tau_{startF}}}\right) \cdot dX$$

**Note**: The boundaries of the integral are inverted to take into account the sign "-" present in the formulation of (L17a).

With (L19), it is got:

$$\Lambda_F(u) = \left(\frac{\Lambda_0 \cdot P_F(u)}{\delta X_T}\right) \cdot \int_{X_{down}}^{X_{up}} \left(1 - e^{-\frac{(X - X_{up})}{u \cdot \tau_{startF}}}\right) \cdot dX$$

Integration gives:

$$\Lambda_F(u) = \left(\frac{\Lambda_0 \cdot P_F(u)}{\delta X_T}\right) \cdot \left[\delta X_{Max} + u \cdot \tau_{startF} \cdot \left(1 - e^{\frac{\delta X_{Max}}{u \cdot \tau_{startF}}}\right)\right] \qquad \text{(L23)}$$

**Tension generated by the $\Lambda_F(u)$ heads in WS according to {startF}**

With (L9a) the equation (L16a) is reformulated:

$$pT_F^*(i\cdot\tau_{sair}) = \left(\frac{P_{SB}^*}{\delta X_T}\right)\cdot\sum_{k=1}^{n_M}\left[\left(1-e^{-\frac{k\cdot\tau_{stair}}{\tau_{startF}}}\right)\cdot\frac{(\delta X_{Max}-(k-1/2)\cdot\delta X_{stair})}{|X_{down}|}\cdot\delta X_{stair}\right]$$

For a hs shortening at constant speed (u), we use (L17a) and (L17b): the factor ($k\cdot\tau_{stair}$) tends towards t; the expression [$(k-1/2)\cdot\delta X_{stair}$] converges on ($X_{up}$-X) according to (L19); the product ($\delta X_{stair}^2$) tends towards $dX^2$, a negligible term because it is of the 2nd order.

The relative tension generated by the heads that initiated a WS rapidly ($pT^*_F$) in repetitive mode tends towards a relative tension ($pT_F$), a function of u, which is calculated using (L22a) according to:

$$pT_F^* \to pT_F(u) = \left(\frac{P_F(u)}{\delta X_T}\right)\cdot\int_{X_{down}}^{X_{up}}\left[\left(1-e^{-\frac{(X-X_{up})}{u\cdot\tau_{startF}}}\right)\cdot\left(\frac{\delta X_{Max}-(X_{up}-X)}{|X_{down}|}\right)\right]\cdot dX$$

After integration:

$$pT_F(u) = \left[\frac{P_F(u)\cdot\delta X_{Max}}{\delta X_T\cdot|X_{down}|}\right]\cdot\left[\frac{\delta X_{Max}}{2}+u\cdot\tau_{startF}+u^2\cdot\frac{\tau_{startF}^2}{\delta X_{Max}}\cdot\left(1-e^{\frac{\delta X_{Max}}{u\cdot\tau_{startF}}}\right)\right] \qquad (L24)$$

#### *L.5.4 Numbers and tensions generated by WS heads initiated according to {startS} when hs is shortened at constant speed u*

**Number of WS heads having initiated with respect to {startS}**

With (L9b), the expression (L13b) is reformulated:

$$\Lambda_S^*(i\cdot\tau_{sair}) = \left(\frac{\Lambda_0\cdot P_{Slow}^*}{\delta X_T}\right)\cdot\sum_{k=(n_{preS}+1)}^{n_M}\left[\left(1-e^{-\frac{k\cdot\tau_{stair}-\tau_{preS}}{\tau_{startS}}}\right)\cdot\delta X_{stair}\right]$$

For a hs shortening at constant speed, expressions (L17a) and (L17b) are imputed and the number of heads ($\Lambda^*_S$) slowly initiating a WS on $\delta X_{Max}$ converges towards a number ($\Lambda_S$), function of u, which is calculated using (L20) and (L22b) according to:

$$\Lambda_S^* \to \Lambda_S(u) = \left(\frac{\Lambda_0\cdot P_S(u)}{\delta X_T}\right)\cdot\int_{X_{down}}^{X_S}\left(1-e^{-\frac{t-\tau_{preS}}{\tau_{startS}}}\right)\cdot dX$$

With (L19) and (L20a), it is obtained:

$$\Lambda_S(u) = \left(\frac{\Lambda_0\cdot P_S(u)}{\delta X_T}\right)\cdot\int_{X_{down}}^{X_S}\left(1-e^{-\frac{(X-X_S)}{u\cdot\tau_{startS}}}\right)\cdot dX$$

After integration:

$$\Lambda_S(u)=\left(\frac{\Lambda_0\cdot P_S(u)}{\delta X_T}\right)\cdot\left[\left(X_S+\left|X_{down}\right|\right)+u\cdot\tau_{startS}\cdot\left(1-e^{\frac{X_S+\left|X_{down}\right|}{u\cdot\tau_{startS}}}\right)\right] \qquad \text{(L25)}$$

**Tension generated by the $\Lambda_S$(u) WS heads according to {startS}**

The relative tension generated by the heads that initiated a WS slowly in repetitive regime ($pT^*_S$) formulated in (L16b) tends towards a relative tension ($pT_S$), function of u, which is calculated by applying (L22b):

$$pT_S(u)=\left(\frac{P_S(u)}{\left|X_{down}\right|\cdot\delta X_T}\right)\cdot\int_{X_{down}}^{X_S}\left[\left(1-e^{-\frac{(X-X_S)}{u\cdot\tau_{startS}}}\right)\cdot\left(\left|X_{down}\right|+X\right)\right]\cdot dX$$

Integration gives:

$$pT_S(u)=\left[\frac{P_S(u)\cdot\left(X_S+\left|X_{down}\right|\right)}{\left|X_{down}\right|\cdot\delta X_T}\right]\cdot\left[\frac{\left(X_S+\left|X_{down}\right|\right)}{2}+u\cdot\tau_{startS}+\frac{u^2\cdot\tau_{startS}{}^2}{\left(X_S+\left|X_{down}\right|\right)}\cdot\left(1-e^{\frac{X_S+\left|X_{down}\right|}{u\cdot\tau_{startS}}}\right)\right]$$

(L26)

##### *L.5.5 Numbers and tensions generated by WS heads initiated according to {startVS} when hs is shortened at constant speed u*

**Number of WS heads having initiated according to {startVS}**

Using the same method, the number of heads ($\Lambda^*_{VS}$) initiating a WS on $\delta X_{Max}$ very slowly converges to a number ($\Lambda_{VS}$) function of u that is calculated using (L20) and (L22c) according to :

$$\Lambda^*_{VS}\rightarrow\Lambda_{VS}(u)=\left(\frac{\Lambda_0\cdot P_{VS}(u)}{\delta X_T}\right)\cdot\int_{X_{down}}^{X_{VS}}\left(1-e^{-\frac{t-\tau_{preVS}}{\tau_{startVS}}}\right)\cdot dX$$

With (L19) and (L20b), we obtain:

$$\Lambda_{VS}(u)=\left(\frac{\Lambda_0\cdot P_{VS}(u)}{\delta X_T}\right)\cdot\int_{X_{down}}^{X_{VS}}\left(1-e^{-\frac{(X-X_{VS})}{u\cdot\tau_{startVS}}}\right)\cdot dX$$

After integration:

$$\Lambda_{VS}(u)=\left(\frac{\Lambda_0\cdot P_{VS}(u)}{\delta X_T}\right)\cdot\left[\left(X_{VS}+\left|X_{down}\right|\right)+u\cdot\tau_{startVS}\cdot\left(1-e^{\frac{X_{VS}+\left|X_{down}\right|}{u\cdot\tau_{startVS}}}\right)\right] \qquad \text{(L27)}$$

**Tension generated by the $\Lambda_{VS}(u)$ WS heads according to {startVS}**

The relative tension generated by the heads that initiated a WS very slowly in repetitive mode ($pT^*_{VS}$) formulated in (16c) tends towards a relative tension (pTVS), function of u, which is calculated by applying (L22c) according to:

$$pT_{VS}(u)=\left(\frac{P_{VS}(u)}{|X_{down}|\cdot\delta X_T}\right)\cdot\int_{X_{down}}^{X_{VS}}\left[\left(1-e^{-\frac{(X-X_{VS})}{u\cdot\tau_{startVS}}}\right)\cdot\left(|X_{down}|+X\right)\right]\cdot dX$$

Integration gives:

$$pT_{VS}(u)=\left[\frac{P_{VS}(u)\cdot\left(X_{VS}+|X_{down}|\right)}{|X_{down}|\cdot\delta X_T}\right]\cdot\left[\frac{\left(X_{VS}+|X_{down}|\right)}{2}+u\cdot\tau_{startVS}+\frac{u^2\cdot\tau_{startVS}{}^2}{\left(X_{VS}+|X_{down}|\right)}\cdot\left(1-e^{\frac{X_{VS}+|X_{down}|}{u\cdot\tau_{startVS}}}\right)\right] \quad \text{(L28)}$$

#### L.6 Calculation of the tension generated by the slowly detaching heads in repetitive regime

##### *L.6.1 Instantaneous proportions of slowly detaching heads in repetitive mode*

Slow detachment is carried out with the {SlowDE} event studied in paragraph B.7 of Supplement S1.B of Paper 1. The {SlowDE} event only occurs for heads with a lever angle θ between $\theta_{up}$ and $\theta_T$ (see empty area in Fig L1a). Thus among the $n_M$ steps of the repetitive regime, only $n_E$ steps are concerned. We pose:

$$n_T = n_M - n_E \quad \text{(L29a)}$$

$$n_{SDE} = n_T + n_{preSDE} \quad \text{(L29b)}$$

where $n_M$, $n_E$ and $n_{preSDE}$ are defined in (L5a), (L5b) and (L7c).

The heads likely to detach slowly are divided into two types according to whether their initiation to the WS state is performed before or after the $\theta_T$ terminal.

**<u>Type 1</u>: WS heads that initiated quickly, slowly and very slowly between $\theta_{up}$ and $\theta_T$**

In repetitive mode, the maximum proportions of WS Type 1 heads are obtained at the step preceding the $\theta_T$ terminal, i.e. step $n_T$ according to (L29a). With (B13), the instantaneous probabilities of slow detachment Type 1 in repetitive mode ($P^*_{F\_SDE1}$ , $P^*_{S\_SDE1}$ and $P^*_{VS\_SDE1}$) for heads that initiated a WS fast, slowly and very slowly between $\theta_{up}$ and $\theta_T$ are determined using (L9a), (L9b) and (L9c), respectively:

$$P^*_{F\_SDE1}(t)=P^*_{SB}\cdot\left(1-e^{-\frac{n_T\cdot\tau_{stair}}{\tau_{startF}}}\right)\cdot\left(1-e^{-\frac{t-\tau_{preSDE}}{\tau_{SDE}}}\right) \quad \text{(L30a)}$$

$$P^*_{S\_SDE1}(t) = P^*_{Slow} \cdot \left(1 - e^{-\frac{(n_T - n_{preS}) \cdot \tau_{stair}}{\tau_{startS}}}\right) \cdot \left(1 - e^{-\frac{t - \tau_{preSDE}}{\tau_{SDE}}}\right) \quad \text{(L30b)}$$

$$P^*_{VS\_SDE1}(t) = P^*_{VSlow} \cdot \left(1 - e^{-\frac{(n_T - n_{preVS}) \cdot \tau_{stair}}{\tau_{startVS}}}\right) \cdot \left(1 - e^{-\frac{t - \tau_{preSDE}}{\tau_{SDE}}}\right) \quad \text{(L30c)}$$

The instantaneous probability of slow detachment in repetitive mode for Type 1 heads ($P^*_{SDE\_1}$) having initiated a WS according to the three modes between $\theta_{up}$ and $\theta_T$ is the sum of the three previous probabilities:

$$P^*_{SDE1}(t) = P^*_{F\_SDE1}(t) + P^*_{S\_SDE1}(t) + P^*_{VS\_SDE1}(t) \quad \text{(L31)}$$

**<u>Type 2</u>: heads that initiate quickly, slowly and very slowly between $\theta_T$ and $\theta_{down}$**

The instantaneous probabilities of slow detachment following the three types of initiation were studied in paragraph B.7 of Supplement S1.B of Paper 1. The maximum proportions of WS Type 2 heads are calculated by the difference between the maximum proportions ($P^*_{SB}$, $P^*_{Slow}$ and $P^*_{VSlow}$) and the maximum proportions calculated for Type 1 heads.

According to (L9a) and (B15a), the instantaneous probability of slow detachment in repetitive mode for Type 2 heads ($P^*_{F\_SDE2}$) having initiated a WS quickly between $\theta_T$ and $\theta_{down}$ is equal to:

$$P^*_{F\_SDE2}(t) = \left(P^*_{SB} \cdot e^{-\frac{n_T \cdot \tau_{stair}}{\tau_{startF}}}\right) \cdot \left(1 - e^{-\frac{t - \tau_{preSDE}}{\tau_{SDE}}}\right) \quad \text{(L32a)}$$

According to (L9b) and (B15b), the instantaneous probability of slow detachment in repetitive mode for Type 2 heads ($P^*_{S\_SDE2}$) having initiated a WS slowly between $\theta_T$ and $\theta_{down}$ is equal to:

$$P^*_{S\_SDE2}(t) = \left(P^*_{Slow} \cdot e^{-\frac{(n_T - n_{preS}) \cdot \tau_{stair}}{\tau_{startS}}}\right) \cdot \left(1 - \frac{\tau_{startVS} \cdot e^{-\frac{t-(\tau_{preS}+\tau_{preSDE})}{\tau_{startVS}}}}{(\tau_{startVS} - \tau_{SDE})} - \frac{\tau_{SDE} \cdot e^{-\frac{t-(\tau_{preS}+\tau_{preSDE})}{\tau_{SDE}}}}{(\tau_{SDE} - \tau_{startVS})}\right)$$

(L32b)

According to (L9c) and (B15c), the instantaneous probability of slow detachment in repetitive mode for Type 2 heads ($P^*_{VS\_SDE2}$) having initiated a WS very slowly between $\theta_T$ and $\theta_{down}$ is equal to:

$$P^*_{VS\_SDE2}(t) = \left(P^*_{VSlow} \cdot e^{-\frac{(n_T - n_{preVS}) \cdot \tau_{stair}}{\tau_{startVS}}}\right) \cdot \left(1 - \frac{\tau_{startVS} \cdot e^{-\frac{t-(\tau_{preVS}+\tau_{preSDE})}{\tau_{startVS}}}}{(\tau_{startVS} - \tau_{SDE})} - \frac{\tau_{SDE} \cdot e^{-\frac{t-(\tau_{preSV}+\tau_{preSDE})}{\tau_{SDE}}}}{(\tau_{SDE} - \tau_{startVS})}\right)$$

(L32c)

The instantaneous probability of slow detachment in repetitive mode for Type 2 heads ($P^*_{SDE\_2}$) having initiated a WS according to the three modes between $\theta_T$ and $\theta_{down}$ is the sum of the three previous probabilities:

$$P^*_{SDE2}(t) = P^*_{F\_SDE2}(t) + P^*_{S\_SDE2}(t) + P^*_{VS\_SDE2}(t) \qquad \text{(L33)}$$

**In total, summing for the two types with (L31) and (L33):**

$$P^*_{SDE}(t) = P^*_{SDE1}(t) + P^*_{SDE2}(t) \qquad \text{(L34)}$$

##### *L.6.2 Number of heads slowly detaching in repetitive regime*

The total number of heads slowly detaching per hs ($\Lambda^*_{SDE}$) at the time end of step n° i in repetitive mode is calculated by summing the expression (L34) for each of the ($n_M$-$n_{SDE}$) stairs and by using the equality (I11) of the Supplement S4.I of Paper 4:

$$\Lambda^*_{SDE}(i \cdot \tau_{sair}) = \left(\Lambda_0 \cdot \frac{\delta X_{stair}}{\delta X_T}\right) \cdot \sum_{k=(n_{preSDE}+1)}^{n_E} \left[P^*_{SDE}(k \cdot \tau_{stair})\right] \qquad \text{(L35)}$$

Equality (L35) is being rewritten:

$$\Lambda^*_{SDE}(i \cdot \tau_{sair}) = \Lambda^*_{F\_SDE1}(i \cdot \tau_{sair}) + \Lambda^*_{S\_SDE1}(i \cdot \tau_{sair}) + \Lambda^*_{VS\_SDE1}(i \cdot \tau_{sair})$$
$$+ \Lambda^*_{F\_SDE2}(i \cdot \tau_{sair}) + \Lambda^*_{S\_SDE2}(i \cdot \tau_{sair}) + \Lambda^*_{VS\_SDE2}(i \cdot \tau_{sair}) \qquad \text{(L36)}$$

where the six expressions of the right member are formulated from (L35) according to (L30a), (L30b), (L30c), (L32a), (L32b) and (L32c), respectively:

$$\Lambda^*_{F\_SDE1}(i \cdot \tau_{sair}) = \left(\Lambda_0 \cdot \frac{\delta X_{stair}}{\delta X_T}\right) \cdot \sum_{k=(n_{preSDE}+1)}^{n_E} \left[P^*_{F\_SDE1}(k \cdot \tau_{stair})\right] \qquad \text{(L37a)}$$

$$\Lambda^*_{S\_SDE1}(i \cdot \tau_{sair}) = \left(\Lambda_0 \cdot \frac{\delta X_{stair}}{\delta X_T}\right) \cdot \sum_{k=(n_{preSDE}+1)}^{n_E} \left[P^*_{S\_SDE1}(k \cdot \tau_{stair})\right] \qquad \text{(L37b)}$$

$$\Lambda^*_{VS\_SDE1}(i \cdot \tau_{sair}) = \left(\Lambda_0 \cdot \frac{\delta X_{stair}}{\delta X_T}\right) \cdot \sum_{k=(n_{preSDE}+1)}^{n_E} \left[P^*_{VS\_SDE1}(k \cdot \tau_{stair})\right] \qquad \text{(L37c)}$$

$$\Lambda^*_{F\_SDE2}(i \cdot \tau_{sair}) = \left(\Lambda_0 \cdot \frac{\delta X_{stair}}{\delta X_T}\right) \cdot \sum_{k=(n_{preSDE}+1)}^{n_E} \left[P^*_{F\_SDE2}(k \cdot \tau_{stair})\right] \qquad \text{(L37d)}$$

$$\Lambda^*_{S\_SDE2}(i \cdot \tau_{sair}) = \left(\Lambda_0 \cdot \frac{\delta X_{stair}}{\delta X_T}\right) \cdot \sum_{k=(n_S+n_{preSDE}+1)}^{n_E} \left[P^*_{S\_SDE2}(k \cdot \tau_{stair})\right] \qquad \text{(L37e)}$$

$$\Lambda^*_{VS\_SDE2}(i \cdot \tau_{sair}) = \left(\Lambda_0 \cdot \frac{\delta X_{stair}}{\delta X_T}\right) \cdot \sum_{k=(n_{VS}+n_{preSDE}+1)}^{n_E} \left[P^*_{VS\_SDE2}(k \cdot \tau_{stair})\right] \qquad \text{(L37f)}$$

##### *L.6.3 Tension created by slowly detaching heads during the repetitive regime*

The calculation for a single step is carried out at the time end of staircase step n° [i-(k-1)] corresponding to the k value.

**Conditions with (L34):**

$$\left.\begin{array}{r} p = P^{*}_{SDE}(k\cdot\tau_{stair}) \\ X_1 = X_T - (k-1)\cdot\delta X_{stair} \\ X_2 = X_T - k\cdot\delta X_{stair} \end{array}\right\} \Rightarrow \begin{array}{c} \delta X_L = X_1 - X_2 = \delta X_{stair} \\ \\ X_1 + X_2 = 2\cdot X_T - (2k-1)\cdot\delta X_{stair} \end{array}$$

**Application with (I31b) in Supplement S4.I to Paper 4:**

$$pT_{SDE}(k\cdot\tau_{stair}) = P^{*}_{SDE}(k\cdot\tau_{stair})\cdot\left(\frac{\delta X_{stair}}{\delta X_T}\right)\cdot\frac{(\delta X_E - (k-1/2)\cdot\delta X_{stair})}{|X_{down}|}$$

By summing the tensions relative to each of the ($n_M$-$n_{SDE}$) stairs, the total relative tension ($pT^*_{SDE}$) generated by the $\Lambda^*_{SDE}$ heads slowly detaching at the time end of step n° i (with i ≥ n*) in repetitive mode is:

$$pT^{*}_{SDE}(i\cdot\tau_{stair}) = \left(\frac{\delta X_{stair}}{\delta X_T}\right)\cdot\sum_{k=(n_{preSDE}+1)}^{n_E}\left[P^{*}_{SDE}(k\cdot\tau_{stair})\cdot\frac{(\delta X_E - (k-1/2)\cdot\delta X_{stair})}{|X_{down}|}\right] \quad \text{(L38)}$$

Equality (A38) is being rewritten:

$$pT^{*}_{SDE\_stair}(i\cdot\tau_{sair}) = pT^{*}_{F\_SDE1}(i\cdot\tau_{sair}) + pT^{*}_{S\_SDE1}(i\cdot\tau_{sair}) + pT^{*}_{VS\_SDE1}(i\cdot\tau_{sair})$$
$$+ pT^{*}_{F\_SDE2}(i\cdot\tau_{sair}) + pT^{*}_{S\_SDE2}(i\cdot\tau_{sair}) + pT^{*}_{VS\_SDE2}(i\cdot\tau_{sair}) \quad \text{(L39)}$$

where the six expressions of the right member are formulated from (L38) according to (L30a), (L30b), (L30c), (L32a), (L32b) and (L32c), respectively:

$$pT^{*}_{F\_SDE1}(i\cdot\tau_{sair}) = \left(\frac{\delta X_{stair}}{\delta X_T}\right)\cdot\left(\sum_{k=(n_{preSDE}+1)}^{n_E}\left[P^{*}_{F\_SDE1}(k\cdot\tau_{stair})\cdot\frac{(\delta X_E - (k-1/2)\cdot\delta X_{stair})}{|X_{down}|}\right]\right) \quad \text{(L40a)}$$

$$pT^{*}_{S\_SDE1}(i\cdot\tau_{sair}) = \left(\frac{\delta X_{stair}}{\delta X_T}\right)\cdot\left(\sum_{k=(n_{preSDE}+1)}^{n_E}\left[P^{*}_{S\_SDE1}(k\cdot\tau_{stair})\cdot\frac{(\delta X_E - (k-1/2)\cdot\delta X_{stair})}{|X_{down}|}\right]\right) \quad \text{(L40b)}$$

$$pT^{*}_{VS\_SDE1}(i\cdot\tau_{sair}) = \left(\frac{\delta X_{stair}}{\delta X_T}\right)\cdot\left(\sum_{k=(n_{preSDE}+1)}^{n_E}\left[P^{*}_{VS\_SDE1}(k\cdot\tau_{stair})\cdot\frac{(\delta X_E - (k-1/2)\cdot\delta X_{stair})}{|X_{down}|}\right]\right) \quad \text{(L40c)}$$

$$pT^{*}_{F\_SDE2}(i\cdot\tau_{sair}) = \left(\frac{\delta X_{stair}}{\delta X_T}\right)\cdot\left(\sum_{k=(n_{preSDE}+1)}^{n_E}\left[P^{*}_{F\_SDE2}(k\cdot\tau_{stair})\cdot\frac{(\delta X_E - (k-1/2)\cdot\delta X_{stair})}{|X_{down}|}\right]\right) \quad \text{(L40d)}$$

$$pT^{*}_{S\_SDE2}(i\cdot\tau_{sair}) = \left(\frac{\delta X_{stair}}{\delta X_T}\right)\cdot\left(\sum_{k=(n_S+n_{preSDE}+1)}^{n_E}\left[P^{*}_{S\_SDE2}(k\cdot\tau_{stair})\cdot\frac{(\delta X_E - (k-1/2)\cdot\delta X_{stair})}{|X_{down}|}\right]\right) \quad \text{(L40e)}$$

$$pT^{*}_{VS\_SDE2}(i\cdot\tau_{sair}) = \left(\frac{\delta X_{stair}}{\delta X_T}\right)\cdot\left(\sum_{k=(n_{VS}+n_{preSDE}+1)}^{n_E}\left[P^{*}_{VS\_SDE2}(k\cdot\tau_{stair})\cdot\frac{(\delta X_E - (k-1/2)\cdot\delta X_{stair})}{|X_{down}|}\right]\right)$$
(L40f)

#### L.7 Calculation of the tension generated by the slowly detaching heads when a hs is shortening at constant speed

We take up the same reasoning developed in paragraph L.5 where the two parameters characterizing a staircase step, the length ($\delta X_{stair}$) and the duration ($\tau_{stair}$), are gradually decreased until they tend towards infinitesimal values according to (L17a) and (L17b), i.e. the staircase shortening is assimilated for each hs to a shortening performed at a constant speed (u) determined in (L18).

From the abscissa $X_T$ which corresponds to X(t=0) for the event {SlowDE}, the abscissa $X_{SDE}$ is defined as a function of u with respect to the occurrence of the {SlowDE} event:

$$X_{SDE} = X_T + u \cdot \tau_{preSDE} \qquad \text{(L41)}$$

We pose relatively to the slowly detaching Type 1 heads:

$$P_{F\_SDE1}(u) = P_F(u) \cdot \left( 1 - e^{\frac{\delta X_T}{u \cdot \tau_{startF}}} \right) \cdot \mathbf{1}_{[\![ |u| \cdot \tau_{preSDE} \le \delta X_E ]\!]}(u) \qquad \text{(L42a)}$$

$$P_{S\_SDE1}(u) = P_S(u) \cdot \left( 1 - e^{\frac{\delta X_T + u \cdot \tau_{preS}}{u \cdot \tau_{startS}}} \right) \cdot \mathbf{1}_{[\![ |u| \cdot \tau_{preS} \le \delta X_T ]\!]}(u) \cdot \mathbf{1}_{[\![ |u| \cdot \tau_{preSDE} \le \delta X_E ]\!]}(u) \qquad \text{(L42b)}$$

$$P_{VS\_SDE1}(u) = \left(1 - p_{startS} - p_{startF}\right) \cdot \left( 1 - e^{\frac{\delta X_T + u \cdot \tau_{preVS}}{u \cdot \tau_{startVS}}} \right) \cdot \mathbf{1}_{[\![ |u| \cdot \tau_{preVS} \le \delta X_T ]\!]}(u) \cdot \mathbf{1}_{[\![ |u| \cdot \tau_{preSDE} \le \delta X_E ]\!]}(u)$$

(L42c)

And relatively to the slowly detaching Type 2 heads:

$$P_{F\_SDE2}(u) = P_F(u) \cdot \left( e^{\frac{\delta X_T}{u \cdot \tau_{startF}}} \right) \cdot \mathbf{1}_{[\![ |u| \cdot \tau_{preSDE} \le \delta X_E ]\!]}(u) \qquad \text{(L42d)}$$

$$P_{S\_SDE2}(u) = P_S(u) \cdot \left( e^{\frac{\delta X_T + u \cdot \tau_{preS}}{u \cdot \tau_{startS}}} \right) \cdot \mathbf{1}_{[\delta X_T < |u| \cdot \tau_{preS} \le \delta X_{Max}]}(u) \cdot \mathbf{1}_{[\![ |u| \cdot \tau_{preSDE} \le \delta X_E ]\!]}(u) \qquad \text{(L42e)}$$

$$P_{VS\_SDE2}(u) = \left(1 - p_{startS} - p_{startF}\right) \cdot \left( e^{\frac{\delta X_T + u \cdot \tau_{preVS}}{u \cdot \tau_{startVS}}} \right) \cdot \mathbf{1}_{[\delta X_T < |u| \cdot \tau_{preVS} \le \delta X_{Max}]}(u) \cdot \mathbf{1}_{[\![ |u| \cdot \tau_{preSDE} \le \delta X_E ]\!]}(u)$$

(L42f)

##### *L.7.1 Number of heads detaching slowly when the hs is shortened at constant speed u*

With (L30a), the expression (L37a) is reformulated:

$$\overset{*}{\Lambda}_{F\_SDE1}(i\cdot\tau_{sair}) = \left(\frac{\Lambda_0\cdot \overset{*}{P}_{SB}}{\delta X_T}\right)\cdot\left(1-e^{-\frac{n_T\cdot\tau_{stair}}{\tau_{startF}}}\right)\cdot\left(\sum_{k=(n_{preSDE}+1)}^{n_E}\left[\left(1-e^{-\frac{k\cdot\tau_{stair}-\tau_{preSDE}}{\tau_{SDE}}}\right)\cdot\delta X_{stair}\right]\right)$$

At constant speed shortening when applying (L17a) and (L17b), the factor ($k\cdot\tau_{stair}$) tends towards t, and the number of Type 1 heads slowly detaching after quickly initiating a WS between $\theta_{up}$ and $\theta_T$ in repetitive mode ($\Lambda^*_{F\_SDE1}$) tends towards a number ($\Lambda_{F\_SDE1}$), function of u, that is determined by applying (L22a), (L41) and (L42a):

$$\overset{*}{\Lambda}_{F\_SDE1} \rightarrow \Lambda_{F\_SDE1}(u) = \left(\frac{\Lambda_0\cdot P_{F\_SDE1}(u)}{\delta X_T}\right)\cdot\int_{X_{down}}^{X_{SDE}}\left(1-e^{-\frac{(X-X_{SDE})}{u\cdot\tau_{SDE}}}\right)\cdot dX$$

After integration:

$$\Lambda_{F\_SDE1}(u) = \left(\frac{\Lambda_0\cdot P_{F\_SDE1}(u)}{\delta X_T}\right)\cdot\left[\left(X_{SDE}+|X_{down}|\right)+u\cdot\tau_{SDE}\cdot\left(1-e^{\frac{X_{SDE}+|X_{down}|}{u\cdot\tau_{SDE}}}\right)\right] \quad \text{(L43a)}$$

Following the same path, the two numbers of Type 1 heads slowly detaching after slowly and very slowly initiating a WS between $\theta_{up}$ and $\theta_T$ in repetitive mode ($\Lambda^*_{S\_SDE1}$ and $\Lambda^*_{VS\_SDE1}$) tend towards two numbers ($\Lambda_{S\_SDE1}$ and $\Lambda_{VS\_SDE1}$), functions of u, which are calculated by applying (L22b), (L22c), (L41), (L42b) and (L42c):

$$\Lambda_{S\_SDE1}(u) = \left(\frac{\Lambda_0\cdot P_{S\_SDE1}(u)}{\delta X_T}\right)\cdot\left[\left(X_{SDE}+|X_{down}|\right)+u\cdot\tau_{SDE}\cdot\left(1-e^{\frac{X_{SDE}+|X_{down}|}{u\cdot\tau_{SDE}}}\right)\right] \quad \text{(L43b)}$$

$$\Lambda_{VS\_SDE1}(u) = \left(\frac{\Lambda_0\cdot P_{VS\_SDE1}(u)}{\delta X_T}\right)\cdot\left[\left(X_{SDE}+|X_{down}|\right)+u\cdot\tau_{SDE}\cdot\left(1-e^{\frac{X_{SDE}+|X_{down}|}{u\cdot\tau_{SDE}}}\right)\right] \quad \text{(L43c)}$$

The three numbers of Type 2 heads slowly detaching after quickly, slowly and very slowly initiating a WS between $\theta_T$ and $\theta_{down}$ in repetitive mode ($\Lambda^*_{F\_SDE2}$, $\Lambda^*_{S\_SDE2}$ and $\Lambda^*_{VS\_SDE2}$) tend towards three numbers ($\Lambda_{F\_SDE2}$, $\Lambda_{S\_SDE2}$ and $\Lambda_{VS\_SDE2}$), functions of u, which are calculated by applying (L22a), (L22b), (L22c), (L41), (L42d), (L42e) and (L42f):

$$\Lambda_{F\_SDE2}(u) = \left(\frac{\Lambda_0 \cdot P_{F\_SDE2}(u)}{\delta X_T}\right) \cdot \left[ \left( x_{SDE} + |x_{down}| \right) + u \cdot \tau_{SDE} \cdot \left( 1 - e^{-\frac{x_{SDE} + |x_{down}|}{u \cdot \tau_{SDE}}} \right) \right] \quad \text{(L43d)}$$

$$\Lambda_{S\_SDE2}(u) = \left(\frac{\Lambda_0 \cdot P_{S\_SDE2}(u)}{\delta X_T}\right) \cdot \left[ \left( x_{SDE} + |x_{down}| \right) + \frac{u \cdot \tau_{startS}^2}{\left(\tau_{startS} - \tau_{SDE}\right)} \cdot \left( 1 - e^{-\frac{x_{SDE} + |x_{down}|}{u \cdot \tau_{startS}}} \right) + \frac{u \cdot \tau_{SDE}^2}{\left(\tau_{SDE} - \tau_{startS}\right)} \cdot \left( 1 - e^{-\frac{x_{SDE} + |x_{down}|}{u \cdot \tau_{SDE}}} \right) \right] \quad \text{(L43e)}$$

$$\Lambda_{VS\_SDE2}(u) = \left(\frac{\Lambda_0 \cdot P_{VS\_SDE2}(u)}{\delta X_T}\right) \cdot \left[ \left( x_{SDE} + |x_{down}| \right) + \frac{u \cdot \tau_{startVS}^2}{\left(\tau_{startVS} - \tau_{SDE}\right)} \cdot \left( 1 - e^{-\frac{x_{SDE} + |x_{down}|}{u \cdot \tau_{startVS}}} \right) + \frac{u \cdot \tau_{SDE}^2}{\left(\tau_{SDE} - \tau_{startVS}\right)} \cdot \left( 1 - e^{-\frac{x_{SDE} + |x_{down}|}{u \cdot \tau_{SDE}}} \right) \right] \quad \text{(L43f)}$$

##### *L.7.2 Tensions generated by WS heads slowly detaching when the hs is shortening at constant velocity*

The three relative tensions generated by the Type 1 heads slowly detaching after having initiated quickly, slowly and very slowly a WS between $\theta_{up}$ and $\theta_T$ in repetitive mode ($pT^*_{F\_SDE1}$, $pT^*_{S\_SDE1}$ and $pT^*_{VS\_SDE1}$) tend towards three relative tensions ($pT_{F\_SDE1}$, $pT_{S\_SDE1}$ and $pT_{VS\_SDE1}$), functions of u, and equal after integration to:

$$pT_{F\_SDE1}(u) = \left[\frac{P_{F\_SDE1}(u) \cdot \left(x_{SDE} + |x_{down}|\right)}{|x_{down}| \cdot \delta X_T}\right] \cdot \left[ \frac{\left(x_{SDE} + |x_{down}|\right)}{2} + u \cdot \tau_{SDE} + \frac{u^2 \cdot \tau_{SDE}^2}{\left(x_{SDE} + |x_{down}|\right)} \cdot \left( 1 - e^{-\frac{x_{SDE} + |x_{down}|}{u \cdot \tau_{SDE}}} \right) \right] \quad \text{(L44a)}$$

$$pT_{S\_SDE1}(u) = \left[\frac{P_{S\_SDE1}(u) \cdot \left(x_{SDE} + |x_{down}|\right)}{|x_{down}| \cdot \delta X_T}\right] \cdot \left[ \frac{\left(x_{SDE} + |x_{down}|\right)}{2} + u \cdot \tau_{SDE} + \frac{u^2 \cdot \tau_{SDE}^2}{\left(x_{SDE} + |x_{down}|\right)} \cdot \left( 1 - e^{-\frac{x_{SDE} + |x_{down}|}{u \cdot \tau_{SDE}}} \right) \right] \quad \text{(L44b)}$$

$$pT_{VS\_SDE1}(u) = \left[\frac{P_{VS\_SDE1}(u) \cdot \left(x_{SDE} + |x_{down}|\right)}{|x_{down}| \cdot \delta X_T}\right] \cdot \left[ \frac{\left(x_{SDE} + |x_{down}|\right)}{2} + u \cdot \tau_{SDE} + \frac{u^2 \cdot \tau_{SDE}^2}{\left(x_{SDE} + |x_{down}|\right)} \cdot \left( 1 - e^{-\frac{x_{SDE} + |x_{down}|}{u \cdot \tau_{SDE}}} \right) \right] \quad \text{(L44c)}$$

The three relative tensions generated by the Type 2 heads slowly detaching after having initiated quickly, slowly and very slowly a WS between $\theta_T$ and $\theta_{down}$ in repetitive mode ($pT^*_{F\_SDE2}$, $pT^*_{S\_SDE21}$ and $pT^*_{VS\_SDE2}$) tend towards three relative tensions ($pT_{F\_SDE2}$, $pT_{S\_SDE2}$ and $pT_{VS\_SDE2}$), functions of u and equal after integration with:

$$pT_{F\_SDE2}(u)=\left[\frac{P_{F\_SDE2}(u)\cdot\left(x_{SDE}+\left|x_{down}\right|\right)}{\left|x_{down}\right|\cdot\delta X_T}\right]\cdot\left[\frac{\left(x_{SDE}+\left|x_{down}\right|\right)}{2}+u\cdot\tau_{SDE}+\frac{u^2\cdot\tau_{SDE}{}^2}{\left(x_{SDE}+\left|x_{down}\right|\right)}\cdot\left(1-e^{\frac{x_{SDE}+\left|x_{down}\right|}{u\cdot\tau_{SDE}}}\right)\right]$$

(L44d)

$$pT_{S\_SDE2}(u)=\left[\frac{P_{S\_SDE2}(u)\cdot\left(x_{SDE}+\left|x_{down}\right|\right)}{\left|x_{down}\right|\cdot\delta x_T}\right]\cdot$$

$$\left[\frac{\left(x_{SDE}+\left|x_{down}\right|\right)}{2}+u\cdot\left(\tau_{SDE}+\tau_{startS}\right)+\frac{u^2}{\left(x_{SDE}+\left|x_{down}\right|\right)}\cdot\left(\frac{\tau_{SDE}{}^3}{\left(\tau_{SDE}-\tau_{startS}\right)}\left(1-e^{\frac{x_{SDE}+[x_{down}[}{u\cdot\tau_{SDE}}}\right)+\frac{\tau_{startS}{}^3}{\left(\tau_{startS}-\tau_{SDE}\right)}\left(1-e^{\frac{x_{SDE}+\left|x_{down}\right|}{u\cdot\tau_{startS}}}\right)\right)\right]$$

(L44e)

$$pT_{VS\_SDE2}(u)=\left[\frac{P_{VS\_SDE2}(u)\cdot\left(x_{SDE}+\left|x_{down}\right|\right)}{\left|x_{down}\right|\cdot\delta x_T}\right]\cdot$$

$$\left[\frac{\left(x_{SDE}+\left|x_{down}\right|\right)}{2}+u\cdot\left(\tau_{SDE}+\tau_{startVS}\right)+\frac{u^2}{\left(x_{SDE}+\left|x_{down}\right|\right)}\left(\frac{\tau_{SDE}{}^3}{\left(\tau_{SDE}-\tau_{startVS}\right)}\left(1-e^{\frac{x_{SDE}+[x_{down}[}{u\cdot\tau_{SDE}}}\right)+\frac{\tau_{startVS}{}^3}{\left(\tau_{startVS}-\tau_{SDE}\right)}\left(1-e^{\frac{x_{SDE}+\left|x_{down}\right|}{u\cdot\tau_{startVS}}}\right)\right)\right]$$

(L44f)

#### L.8 Total with the 4 events {startF}, {startS}, {startVS} and {SlowDE}

##### *L.8.1 Proportions*

As a reminder, the proportions of the heads initiating a WS when the hs is shortened to constant speed (u) are declined from (L22a), (L22b) and (L22c) according to the 3 respective modes, fast, slow and very slow :

$$P_F(u) = p_{startF} \cdot \left(1 - e^{\frac{\delta X_{pre} + u \cdot \tau_{preSB}}{u \cdot \tau_{SB}}}\right)$$

$$P_S(u) = \left[p_{startS} + p_{startF} - P_F(u)\right] \cdot \mathbf{1}_{\left[|u| \cdot \tau_{preS} \leq \delta X_{Max}\right]}(u)$$

$$P_{VS}(u) = \left(1 - p_{startF} - p_{startS}\right) \cdot \mathbf{1}_{\left[|u| \cdot \tau_{preVS} \leq \delta X_{Max}\right]}(u)$$

The proportions of the slowly detaching heads are classified into 2 types for each of the 3 initiation modes of a WS, i.e. 6 proportions established respectively with equations (L42a) to (L42f). As this event only concerns very slow speeds, we neglect the proportions of the Type 2 heads to simplification purposes, that is by summing (L42a), (L42b) and (L42c):

$$P_{SDE}(u) = P_{SDE1\_F}(u) + P_{SDE1\_S}(u) + P_{SDE1\_VS}(u)$$

with $P_{SDE2\_F}(u) \approx 0$

$P_{SDE2\_S}(u) \approx 0$

$P_{SDE2\_VS}(u) \approx 0$

So:

$$P_{SDE}(u) = \left[P_F(u) \cdot \left(1 - e^{\frac{\delta X_T}{u \cdot \tau_{startF}}}\right) + P_S(u) \cdot \left(1 - e^{\frac{\delta X_T + u \cdot \tau_{preS}}{u \cdot \tau_{startS}}}\right) + P_{VS}(u) \cdot \left(1 - e^{\frac{\delta X_T + u \cdot \tau_{preVS}}{u \cdot \tau_{startVS}}}\right)\right] \cdot \mathbf{1}_{\left[|u| \cdot \tau_{preSDE} \leq \delta X_E\right]}(u)$$

(L45)

***Note: The proportions relative to the slowly detaching heads in Type 2 are negligible and therefore neglected in order to simplify the writing of the equations leading to the Force-Speed relationship but are integrated into our calculations for accuracy, particularly in the algorithms leading to the plots of the F-V relationship of Paper 1.***

##### *L.8.2 Numbers*

By definition, the number of heads when hs is shortened at constant speed u is equal to:

$$\Lambda(u) = \Lambda_F(u) + \Lambda_S(u) + \Lambda_{VS}(u) - \Lambda_{SDE}(u)$$

With (L23), (L25), (L27), (L43a), (L43b), (L43c) and (L45):

$$\begin{aligned}
\Lambda(u) = &\left(\frac{\Lambda 0 \cdot P_F(u)}{\delta X_T}\right)\cdot\left[\delta X_{Max} + u\cdot\tau_{startF}\cdot\left(1-e^{\frac{\delta X_{Max}}{u\cdot\tau_{startF}}}\right)\right]\\
&+\left(\frac{\Lambda 0 \cdot P_S(u)}{\delta X_T}\right)\cdot\left[\left(X_S+|X_{down}|\right)+u\cdot\tau_{startS}\cdot\left(1-e^{\frac{X_S+|X_{down}|}{u\cdot\tau_{startS}}}\right)\right]\\
&+\left(\frac{\Lambda 0 \cdot P_{VS}(u)}{\delta X_T}\right)\cdot\left[\left(X_{VS}+|X_{down}|\right)+u\cdot\tau_{startVS}\cdot\left(1-e^{\frac{X_{VS}+|X_{down}|}{u\cdot\tau_{startVS}}}\right)\right]\\
&-\left(\frac{\Lambda 0 \cdot P_{SDE}(u)}{\delta X_T}\right)\cdot\left[\left(X_{SDE}+|X_{down}|\right)+u\cdot\tau_{SDE}\left(1-e^{\frac{X_{SDE}+|X_{down}|}{u\cdot\tau_{SDE}}}\right)\right]
\end{aligned}\tag{L46}$$

where $X_S = (X_{up} + u\cdot\tau_{preS})$

$X_{VS} = (X_{up} + u\cdot\tau_{preVS})$

$X_{SDE} = (X_T + u\cdot\tau_{preSDE})$

##### *L.8.3 Tensions*

By definition, the relative tension when hs is shortening at constant speed u is equal to :

$$pT(u) = pT_F(u) + pT_S(u) + pT_{VS}(u) - pT_{SDE}(u)$$

So with (L24), (L26), (L28), (L44a), (L44b), (L44c) and (L45):

$$\begin{aligned}
pT(u) = &\left[\frac{P_F(u)\cdot\delta X_{Max}}{|X_{down}|\cdot\delta X_T}\right]\cdot\left[\frac{\delta X_{Max}}{2}+u\cdot\tau_{startF}+u^2\cdot\frac{\tau_{startF}{}^2}{\delta X_{Max}}\cdot\left(1-e^{\frac{\delta X_{Max}}{u\cdot\tau_{startF}}}\right)\right]\\
&+\left[\frac{P_S(u)\cdot\left(X_S+|X_{down}|\right)}{|X_{down}|\cdot\delta X_T}\right]\cdot\left[\frac{\left(X_S+|X_{down}|\right)}{2}+u\cdot\tau_{startS}+\frac{u^2\cdot\tau_{startS}{}^2}{\left(X_{VS}+|X_{down}|\right)}\cdot\left(1-e^{\frac{X_S+|X_{down}|}{u\cdot\tau_{startS}}}\right)\right]\\
&+\left[\frac{P_{VS}(u)\cdot\left(X_{VS}+|X_{down}|\right)}{|X_{down}|\cdot\delta X_T}\right]\cdot\left[\frac{\left(X_{VS}+|X_{down}|\right)}{2}+u\cdot\tau_{startVS}+\frac{u^2\cdot\tau_{startVS}{}^2}{\left(X_{VS}+|X_{down}|\right)}\cdot\left(1-e^{\frac{X_{VS}+|X_{down}|}{u\cdot\tau_{startVS}}}\right)\right]\\
&-\left[\frac{P_{SDE}(u)\cdot\left(X_{SDE}+|X_{down}|\right)}{|X_{down}|\cdot\delta X_T}\right]\cdot\left[\frac{\left(X_{SDE}+|X_{down}|\right)}{2}+u\cdot\tau_{SDE}+\frac{u^2\cdot\tau_{SDE}{}^2}{\left(X_{SDE}+|X_{down}|\right)}\cdot\left(1-e^{\frac{X_{SDE}+|X_{down}|}{u\cdot\tau_{SDE}}}\right)\right]
\end{aligned}\tag{L47}$$

##### *L.8.4 Remarks*

In all previous formulations, the velocity u is expressed as algebraic value and is therefore negative for a hs shortening. Conventionally in the articles, u represents the positive modulus of shortening velocity; also by adding a sign "-", to equations (L45), (L46) and (L47), we find the formulas given previously in accompanying Paper 1.

#### L.9 Density of the angle θ as a function of the shortening speed (u)

##### *L.9.1 Linear relationship between time t and angle θ*

Time (t) is an affine function of the angle θ according to the equality (L19) transformed with (L1):

$$t = \frac{L_{S1b} \cdot R_{WS} \cdot \left(\theta - \theta_{up}\right)}{u} \tag{L48}$$

where u is the shortening velocity common to all hs of the fiber; u is expressed in algebraic value, i.e. $u < 0$.

##### *L.9.2 Event {WSstart}*

The expression (L22a) which gives the probability ($P_F$) of occurrence of the event {startF} as a function of the hs shortening velocity (u) is rewritten using (L1) as:

$$P_F(u) = p_{startF} \cdot \left( 1 - e^{\frac{L_{S1b} \cdot R_{WS} \cdot \delta\theta_{pre} + u \cdot \tau_{preSB}}{u \cdot \tau_{SB}}} \right) \tag{L49a}$$

The expressions (L22b) and (L22c) that provide the probabilities ($P_S$ and $P_{VS}$) of realisation of the {startS} and {startVS} events become:

$$P_S(u) = \left[p_{startS} + p_{startF} - P_F(u)\right] \cdot \mathbf{1}_{\left\{\left|\theta_S - \theta_{up}\right| \le \delta\theta_{Max}\right\}}(\theta)(u) \tag{L49b}$$

$$P_{VS}(u) = \left[1 - p_{startF} - p_{startS}\right] \cdot \mathbf{1}_{\left\{\left|\theta_{VS} - \theta_{up}\right| \le \delta\theta_{Max}\right\}}(\theta) \tag{L49c}$$

where $\theta_S$ and $\theta_{VS}$ are the angles obtained after affine transformation of (L20a) and (L20b) with (L1):

$$\theta_S = \theta_{up} + \frac{u \cdot \tau_{preS}}{L_{S1b} \cdot R_{WS}} \tag{L50}$$

$$\theta_{VS} = \theta_{up} + \frac{u \cdot \tau_{preVS}}{L_{S1b} \cdot R_{WS}} \tag{L51}$$

In support of the relationships (B5a), (B5b) and (B5c) presented in Supplement S1.B to Paper 1 and the expressions (L48) to (L51), the densities ($f_F$, $f_S$ and $f_{VS}$) function of θ, the angle of the levers belonging to the heads that initiated a WS quickly, slowly or very slowly, are formulated:

$$f_F(\theta,u) = \left(\frac{P_F(u)}{\delta\theta_T}\right)\cdot\left(1-e^{\frac{L_{S1b}\cdot R_{WS}\cdot(\theta_{up}-\theta)}{u\cdot\tau_{startF}}}\right) \quad \text{(L52a)}$$

$$f_S(\theta,u) = \left(\frac{P_S(u)}{\delta\theta_T}\right)\cdot\left(1-e^{\frac{L_{S1b}\cdot R_{WS}\cdot(\theta_S-\theta)}{u\cdot\tau_{startS}}}\right)\cdot\mathbf{1}_{\{|\theta_S-\theta_{up}|\leq\delta\theta_{Max}\}}(\theta) \quad \text{(L52b)}$$

$$f_{VS}(\theta,u) = \left(\frac{P_{VS}(u)}{\delta\theta_T}\right)\cdot\left(1-e^{\frac{L_{S1b}\cdot R_{WS}\cdot(\theta_{VS}-\theta)}{u\cdot\tau_{startVS}}}\right)\cdot\mathbf{1}_{\{|\theta_{VS}-\theta_{up}|\leq\delta\theta_{Max}\}}(\theta) \quad \text{(L52c)}$$

The three densities are reported to the range $\delta\theta_T$ in accordance with the equality (I.11) of Supplement S4.I of Paper 4.

##### *L.9.3 Event {SlowDE}*

A myosin head slowly detaches according to the event {SlowDE}. In paragraph L.7, the principles of infinitesimal calculation applied to {SlowDE} provide six proportions of slowly detaching heads with equations (L42a) to (L42f). As the three components relating to Type 2 are negligible, they are not taken into account. Expressions (L22a), (L22b) and (L22c) associated with the probability of realisation of the {SlowDE} event defined in (B13) in Supplement S1.B of Paper 1, the density ($f_{SDE}$) is derived from the angle θ of the levers belonging to the slow detaching heads:

$$f_{SDE}(\theta,u) = \left[\frac{P_{F\_SDE1}(u)+P_{S\_SDE1}(u)+P_{VS\_SDE1}(u)}{\delta\theta_T}\right]\cdot\left(1-e^{\frac{L_{S1b}\cdot R_{WS}\cdot(\theta_{SDE}-\theta)}{u\cdot\tau_{SDE}}}\right) \quad \text{(L53)}$$

with $$P_{F\_SDE1}(u) = P_F(u)\cdot\left(1-e^{\frac{L_{S1b}\cdot R_{WS}\cdot\delta\theta_T}{u\cdot\tau_{startF}}}\right)\cdot\mathbf{1}_{\{|\theta_{SDE}-\theta_T|\leq\delta\theta_E\}}(\theta) \quad \text{(L54a)}$$

$$P_{S\_SDE1}(u) = P_S(u)\cdot\left(1-e^{\frac{L_{S1b}\cdot R_{WS}\cdot\delta\theta_T+u\cdot\tau_{preS}}{u\cdot\tau_{startS}}}\right)\cdot\mathbf{1}_{\{|\theta_S-\theta_{up}|\leq\delta\theta_T\}}(\theta)\cdot\mathbf{1}_{\{|\theta_{SDE}-\theta_T|\leq\delta\theta_E\}}(\theta) \quad \text{(L54b)}$$

$$P_{VS\_SDE1}(u) = P_{VS}(u) \cdot \left(1 - e^{-\frac{L_{S1b} \cdot R_{WS} \cdot \delta\theta_T + u \cdot \tau_{preVS}}{u \cdot \tau_{startVS}}}\right) \cdot \mathbf{1}_{\{|\theta_{VS} - \theta_{up}| \leq \delta\theta_T\}}(\theta) \cdot \mathbf{1}_{\{|\theta_{SDE} - \theta_T| \leq \delta\theta_E\}}(\theta)$$

(L54c)

Where $\theta_{SDE}$ is the angle obtained by affine transformation of equality (L41) according to (L1):

$$\theta_{SDE} = \theta_T + \frac{u \cdot \tau_{preSDE}}{L_{S1b} \cdot R_{WS}} \quad \text{(L55)}$$

The density is referred to the angular range $\delta\theta_T$ in accordance with the equality (I.11) of Supplement S4.I of Paper 4.

##### *L.9.4 Expression of the density of the angle θ as a function of u*

The density (f) of the angle θ of the levers belonging to the heads actually in WS during shortening at constant speed is the sum of the three densities established in (L52a), (L52b) and (L52c) relating to the WS heads initiated in one of the three modes, minus the density defined in (L53) relating to the slowly detaching heads, that is:

$$f(\theta,u) = f_F(\theta,u) + f_S(\theta,u) + f_{VS}(\theta,u) - f_{SDE}(\theta,u)$$
